# Differentially abundant bacteria drive the N_2_-fixation of a widespread moss in the forest-tundra transition zone

**DOI:** 10.1101/2023.06.16.545342

**Authors:** Dennis Alejandro Escolástico-Ortiz, Charlotte Blasi, Jean-Philippe Bellenger, Nicolas Derome, Juan Carlos Villarreal-A

**Affiliations:** Département de Biologie, Université Laval, Québec, G1V 0A6, Canada; Institut de Biologie Intégrative et des Systèmes (IBIS), Université Laval, Québec, G1V 0A6, Canada; Centre d’études nordiques (CEN), Université Laval, Québec, QC, Canada; Centre Sève, Département de Chimie, Université de Sherbrooke, Sherbrooke, J1K 2R1, QC, Canada

**Keywords:** *Azorhizobium*, biological nitrogen fixation, core microbiome, moss symbiosis, *Racomitrium lanuginosum*, *Rhodomicrobium*.

## Abstract

Bryophytes maintain symbiosis with epiphytic bacteria influencing the local nutrient budget. Moss bacterial communities are composed of a core microbiome and bacteria recruited from environmental sources. Notably, symbiotic N_2_-fixing bacteria contribute to the N budget in northern ecosystems through biological nitrogen fixation. This process may be affected by the abundance of diazotrophs and moss nutrient content. We used the abundant moss *Racomitrium lanuginosum* in a forest tundra and shrub tundra in Northern Quebec, Canada, to investigate the bacterial and diazotrophic communities associated with habitat type using amplicon sequencing of the bacterial 16S rRNA and *nifH* genes and test whether the moss core microbiome has recruitment from the soil bacteria community. The *nifH* amplicons and element analysis were used to test the effect of diazotrophic abundance and moss nutrient content on N_2_-fixation activity estimated by acetylene reduction assays. Moss microbial communities between tundra types hosted similar bacterial diversity but differentially abundant groups. The core microbiome of *R. lanuginosum* is composed of bacteria strongly associated with northern mosses with no significant recruitment from the soil. The relative abundances of dominant diazotrophs are significantly correlated with acetylene reduction rates. In contrast, the moss nutrient content did not significantly drive N_2_-fixation. The proteobacterial genera *Azorhizobium* and *Rhodomicrobium* represent newly reported bacteria associated with N_2_-fixation rates in the tundra. We identified critical bacterial groups related to moss-bacterial symbiosis and N_2_-fixation in the forest-tundra transition zone, a changing environment susceptible to climate warming.

## INTRODUCTION

Bryophytes are conspicuous and critical elements in northern ecosystems (Chapin III et al. 1992; Daniëls et al. 2013; Choudhary et al. 2016) and maintain symbiotic associations with several epiphytic bacteria that play essential roles in the environment (Warshan et al. 2017; Holland-Moritz et al. 2018). Bryophyte-bacterial communities seem to be shaped by the host species. For example, in boreal forests, some bacterial groups are frequent and abundant on feather and peat mosses suggesting a species-specific relationship (Bragina et al. 2015; Kostka et al. 2016; Jean et al. 2020; Holland-Moritz et al. 2021; Renaudin et al. 2022b). The environment also affects the bacterial composition of host species and may account for community differences between habitats (Bouchard et al. 2020; Jean et al. 2020; RodríguezLJRodríguez et al. 2022). These bacterial community variations are crucial to understanding how environmental changes may affect the abundance of key groups and would help to predict potential shifts in ecosystem dynamics (Alvarenga and Rousk 2021; Klarenberg et al. 2021).

As in other plants, bryophyte bacterial communities comprise taxa recruited from environmental sources, such as surrounding soil, water currents or air and they form a common bacterial consortium that may perform essential functions in a determined habitat or a core microbiome. Core bacteria could be recognized as abundant taxa occurring in most of the individuals in a determined environment (50-95% prevalence) and are considered characteristic functional members (Shade and Handelsman 2012). For mosses, the core microbiome seems to be composed of taxa strongly associated with bryophyte species (Bragina et al. 2015; Holland-Moritz et al. 2021). However, there is limited information considering the contribution of the environment to the moss core microbiome. Some data show that the microbial communities of mosses and underlying soil contrast in composition (Alcaraz et al. 2018; Klarenberg et al. 2023), but no explicit analyses focus on shared core microbiomes. Identifying the recruitment sources for moss core microbiomes will help understand how plants acquire and maintain essential microbiota through the life cycle.

Among moss-associated bacteria, the N_2_-fixing bacteria or diazotrophs are interesting because they use the nitrogenase enzyme to reduce atmospheric N into bioavailable ammonia (NH_3_) through biological nitrogen fixation (BNF). BNF by moss-bacteria symbioses provides around 1.5 – 2.0 kg N ha^-1^ yr^-1^ in boreal forests and up to 5 kg N ha^-1^ yr^-1^ in arctic tundra (DeLuca et al. 2002; Rousk et al. 2013, 2017b). Notably, moss-cyanobacteria symbiosis is a well-studied interaction where the plant attracts cyanobacteria and can influence colonization and N_2_-fixation (Bay et al. 2013a; Warshan et al. 2017). Many studies have focused on the N_2_-fixation associated with the abundant moss species *Pleurozium schreberi* (Willd. ex Brid.) Mitt., *Hylocomium splendens* (Hedw.) Schimp., *Ptilium crista-castrensis* (Hedw.) De Not. and *Sphagnum* spp. demonstrating their high environmental N_2_ contribution in boreal forests and moderately in the tundra (see Rousk et al. 2013; Carella and Schornack 2018; Jean et al. 2020, 2021). Nonetheless, acrocarpous or erect-growing mosses also provide N_2_ input into northern ecosystems, including *Racomitrium* spp., *Dicranum* spp., *Aulacomnium* spp. and *Polytrichum* spp., but the information is almost restricted to boreal zones with scatter information from arctic environments (Calabria et al. 2020; Stuart et al. 2021; Holland-Moritz et al. 2021; Rzepczynska et al. 2022; Klarenberg et al. 2023). These acrocarpic mosses are particularly abundant in the forest-tundra ecotone compared to feather mosses and sphagnum spp., forming large mats (Gordon et al. 2001; Cutler 2011) and, thus, may represent an essential entry of N_2_ through their associated diazotrophic bacteria. For example, the genus *Racomitrium* is widely distributed in the northern hemisphere from North America to arctic and sub-artic Eurasia (Vitt and Marsh 1988; Larraín et al. 2013; Stech et al. 2013; Hedenäs 2020; Escolástico-Ortiz et al. 2022), covering up to 20 million km^2^ in Europe (Hodgetts 2019). Reports on N_2_-fixation rates indicate that *Racomitrium* spp. have one of the highest N_2_-fixation rates among the acrocarpous mosses in temperate grasslands of Washington (Calabria et al. 2020) and Alaska uplands (Stuart et al. 2021). The BNF of those less-studied moss groups could contribute significantly to N cycling in cold settings.

Environmental and biological variables influence the BNF of moss-bacterial associations (Gentili et al. 2005; Rousk et al. 2017a). Indeed, moss host species and habitat type shape bacterial communities, which ultimately influence N_2_-fixation rates (DeLuca et al. 2002; Bay et al. 2013b; Leppänen et al. 2013; Zhang et al. 2016; Calabria et al. 2020). Nonetheless, the abundance of moss diazotrophic bacteria is also an essential estimator of N_2_-fixation activity due to their intrinsic ability to fix nitrogen. For example, the abundance of the cyanobacterial genus *Nostoc* is positively associated with N_2_-fixation in moss-bacterial symbioses (Calabria et al. 2020; Jean et al. 2020; Alvarenga and Rousk 2021; Holland-Moritz et al. 2021; Renaudin et al. 2022a); even though some lineages may be considered “cheaters” due to their high abundance but a low expression for the nitrogenase-related *nifH* gene (Warshan et al. 2016; Carrell et al. 2019). Other reported bacteria potentially correlated with N_2_-fixation in mosses comprise the cyanobacterial genera *Stigonema*, *Nodullaria*, *Hassallia*, *Mastigocladus* and the proteobacterial groups Methylocystaceae, *Bradyrhizobium, Methylibium* and *Acidisoma* (Leppänen et al. 2013; Holland-Moritz et al. 2021; Renaudin et al. 2022b). Especially, non-cyanobacterial diazotrophs may be important N_2_-fixers in arctic environments where cyanobacteria do not meet their temperature optima for BNF (Gentili et al. 2005). Exploring the diazotroph community could help to detect other critical bacteria mediating the arctic N input.

In addition to diazotroph abundance, nutrients such as phosphorus (P), and nitrogenase metal co-factors, molybdenum (Mo), iron (Fe) and vanadium (V) may influence moss-related N_2_-fixation. In the case of P, the high energetic demand of the nitrogenase enzyme requires those elements to perform well. For boreal feather mosses, P and Mo content enhance the cyanobacterial biomass, and the latter is positively associated with N_2_-fixation rates (Rousk et al. 2017a; Renaudin et al. 2022a). In addition, V is a co-factor of the complementary V-nitrogenase enzyme triggered in Mo-limited conditions and accounts for up to 50% of the N_2_-fixation by cryptogams in cold environments (Zhang et al. 2016; Darnajoux et al. 2019; Bellenger et al. 2020; Harwood 2020; Villarreal A. et al. 2021). This switch from Mo to V-nase activity is exemplified when analyzing Mo:V ratios at a regional scale in Eastern Canada, resulting in a positive relationship with moss-related N_2_-fixation ability in Mo-depleted and colder environments (Renaudin et al. 2022a). Although the nutrient effect on BNF is detectable at large spatial scales (latitudinal gradient), at a local scale, BNF seems to vary according to habitat type and could help predict N contribution in localized areas.

We used the northern moss *Racomitrium lanuginosum* (Hedw.) Brid. as a model to investigate the microbial community composition and N_2_-fixation rates in a forest tundra and shrub tundra in Northern Quebec, Canada. Specifically, we aimed to (1) characterize the bacterial and diazotrophic communities associated with the moss and habitat type; (2) determine the moss core microbiome and core bacterial recruitment from the soil to the moss; (3) evaluate the effect of diazotrophic abundance and moss nutrient content (P, Mo, V, Fe) on moss-microbiome N_2_-fixation rates. Based on previous research, we hypothesized that the microbial composition in *R. lanuginosum* responds to habitat type, the core microbiome has recruitment from the surrounding soil, and N_2_-fixation is significantly affected by the abundance of dominant diazotrophs and nutrient content, mainly Mo. To accomplish the research objectives and test these hypotheses, we used a combination of amplicon sequencing approach (16S rRNA gene for all bacteria and *nifH* gene for diazotrophs) coupled with N_2_-fixation estimates by acetylene reduction assays and moss element analyses in moss populations of a forest-tundra ecotone.

## METHODS

### Study sites and sampling

Samples of the moss *R. lanuginosum* were collected from two different populations located in north-western Hudson Bay in July 2019 (Fig. 1). The first sampling site was Kuujjuarapik (55°16’30 “N; 077°45’30 “W), with a mean annual temperature of -3.6 °C, mean annual precipitation of 640 mm and characterized by subarctic forest-tundra vegetation (Prairie Climate Centre and University of Winnipeg 2022). The forest tundra is considered an ecotone between the boreal forest and the arctic zone, usually identified by the northern boundary matching the tree line, and it is composed of shrubby heathland and forest patches with small-height trees (Payette 1993; Payette et al. 2001). The second population occurs further north in Umiujaq (56°32’N; 76°33’W) with a mean annual temperature of -4.6 °C and mean annual precipitation of 543 mm and dominated by subarctic tundra, which is composed of herbaceous species, grasses, mosses and lichens with a canopy generally reaching less than two meters height (Hare and Ritchie 1972; Payette et al. 2001).

**Fig. 1.**
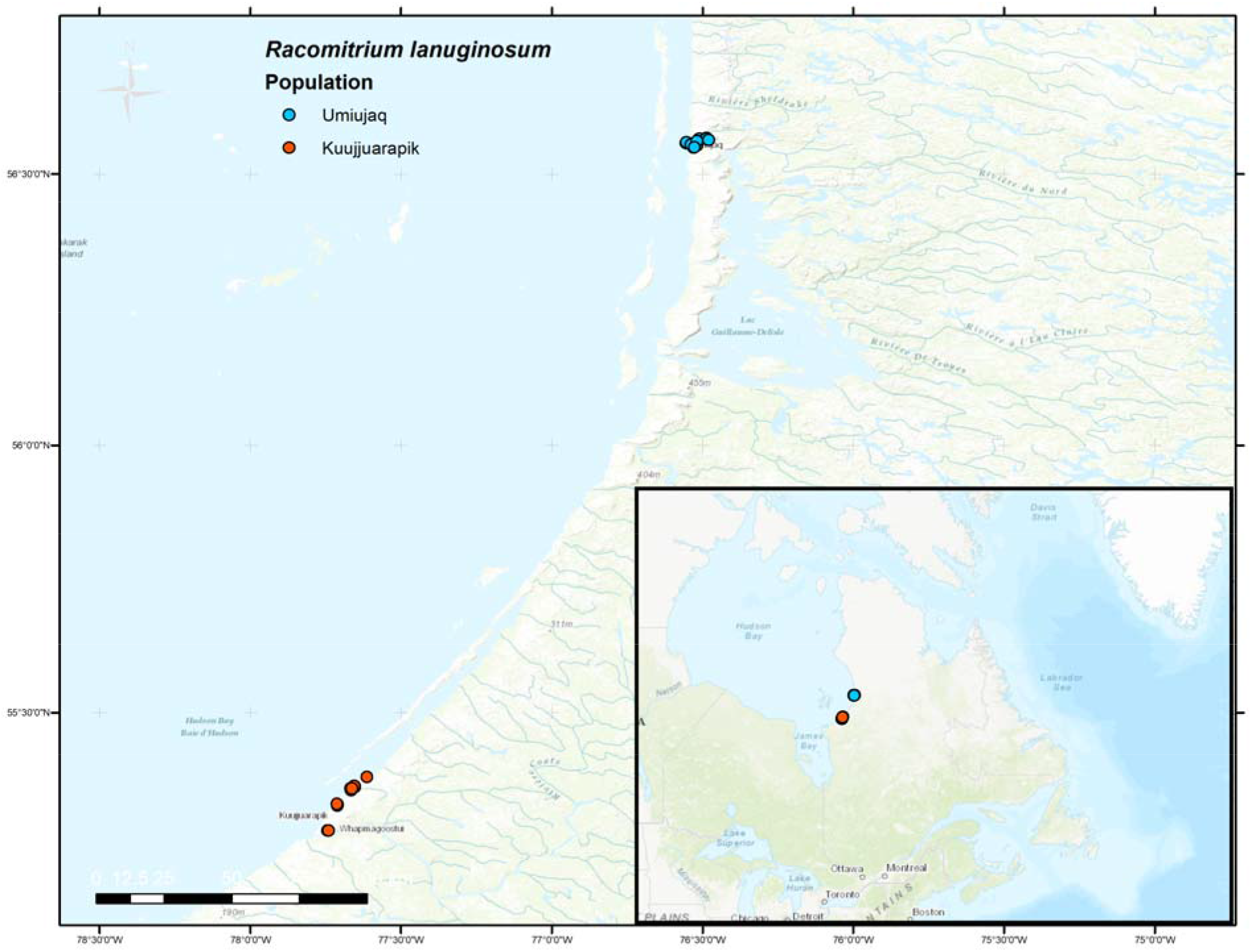
Sampling sites of *Racomitrium lanuginosum* included in this study. The inset map indicates the location of the two populations in the Quebec province, Eastern Canada, including forest tundra in Kuujjuarapik and shrub tundra in Umiujaq

The sampling design consisted of 15 × 15 cm plots with a 3 cm grid (25 squares). We randomly set up 13 plots separated at least by 20 m in each site, Kuujjuarapik and Umiujaq (Fig. S1). In each plot, three squares at an equal distance were selected to control for plot variation. All moss shoots within each square were put in microtubes and selected for microbial analyses. To compare the microbial community of the moss and associated soil, we collected three soil samples from each established plot. All samples were kept in a cooler until arriving at the laboratory. Samples were frozen with liquid nitrogen and homogenized using metal beads. Parallelly, the center of each plot was collected to study the moss microbiome N_2_-fixation ability and moss nutrient content. Vouchers were deposited in the QFA herbarium (Table S1).

### Amplicon sequencing

Moss DNA was isolated from an entire single shoot per sample using the CTAB protocol (Murray and Thompson 1980). DNA extraction of soil samples was performed using the NucleoSpin kit (MACHEREY-NAGEL, Düren, Germany) following the manufacturer’s protocol. We pooled three soil DNA extractions into one mix per plot at an equimolar ratio (Bell-Doyon et al. 2022). The V3-V4 hypervariable regions of the 16S rRNA gene were amplified for moss and soil DNA and the *nifH* gene only for moss DNA to analyze the bacterial community composition. The DNA amplifications consisted of a dual-indexed PCR designed for Illumina instruments by the *Plateforme d’Analyses Génomiques* at the *Institut de Biologie Intégrative et des Systèmes* (IBIS) of *Université Laval* (Quebec City, Canada). The primers for both genomic regions contained Illumina adaptors on the 5’ end: forward-specific primer ACACTCTTTCCCTACACGACGCTCTTCCGATCT-Primer.

The first PCR was performed using the following primers: For the V3-V4 hypervariable regions of the 16S rRNA gene, the forward primer 347R (GGAGGCAGCAGTRRGGAAT) and the reverse 803R (CTACCRGGGTATCTAATCC); for the *nifH* gene, the forward primer Ueda19F (GCI WTY TAY GGI AAR GGI GG) and the reverse primer R6 (GCC ATC ATY TCI CCI GA). The oligonucleotide sequences used for amplification included the selected primers and specific sample indexes conceived for Illumina sequencing.

Amplification of the V3-V4 hypervariable regions of the 16S rRNA gene was carried out using the Q5 high-fidelity DNA polymerase (New England Biolabs, Ipswich, USA). The PCRs were performed in 50 µl reactions. PCR program corresponds to 2 min 98 °C, followed by 35 cycles of 10 s 98 °C, 30 s 55 °C, 30 s 72 °C, and 2 min 72 °C.

To amplify the *nifH* gene, PCRs were done using the TopTaq DNA Polymerase (Qiagen, Venlo, The Netherlands) polymerase in 25 µL reactions. PCR program for *nifH* was 3 min 94 °C, followed by 35 cycles of 30 s 94 °C, 30 s 52 °C, 60 s 72 °C, and 10 min 72 °C. In both cases, we included negative controls to evaluate potential contamination. All PCR products were purified with magnetic beads using 30 µL on 25 µL PCR product for 16S rRNA gene and 35 µL on 25 µL for the *nifH* gene. Amplification, equimolar pooling and sequencing was performed in an Illumina MiSeq technology by the *Plateforme d’analyses génomiques* (IBIS, *Université Laval*, Quebec City, Canada).

### Amplicon sequence data processing

#### *16S* rRNA gene

We used the DADA2 v.1.21 (Callahan et al. 2016) pipeline to process raw sequence data for the 16S rRNA gene. This pipeline infers amplicon sequence variants (ASVs), which provide better resolution, accuracy and reproducibility than other methods and are not strictly constrained by a reference database (Callahan et al. 2017). Each sequencing run was analyzed separately until the ASV inference. Sequences were quality-checked, filtered, the first ten nucleotides removed (*trimLeft*=10) and trimmed with a truncation length of 280 bp for the forward and 230 for the reverse complement (*truncLen*). Amplicons sequence variants (ASVs) were inferred using the default parameters by estimating error rates independently for each run, pooling information across samples (pool= *pseudo*) and merging paired reads to obtain sequence tables per run. Then, we merged sequence tables for the three runs and removed chimeras and sequences with a lower expected length (<400 bp), followed by a taxonomy assignment against the Silva database v.132 (Callahan 2018). The ASV inference analysis recovered 5158 ASVs in 83 samples, including negative controls. The phyloseq package v.1.38 (McMurdie and Holmes 2013) was used to transform the sequence table into a phyloseq object to process the ASV data. First, we excluded chloroplast, mitochondria, phyla with fewer than ten counts and samples with no ASVs. Then, we estimated the library size per sample (Fig. S2) and removed potential contaminants using the decontam v.1.14 package (Davis et al. 2018) using the “frequency” and “prevalence” methods independently. Finally, the ASV’s prevalence was computed to select those in at least 5% of the samples (Figs. S3 and S4). The dataset without the negative controls consisted of 1295 ASVs and 62 samples. The moss and soil samples were treated as subsets into independent phyloseq objects for subsequent analyses.

#### *nifH* gene

Raw sequences of the *nifH* gene were processed in NifMAP v.1.2 (Angel et al. 2018) pipeline. We employed NifMAP because it is specifically designed to manage *nifH* gene data by filtering out homologous sequences with a Hidden-Markov-Model and using a multi-approach method to robustly classify sequences into Operational Taxonomic Units (OTUs) and *nifH* phylogenetic clusters. Before sequence processing, we used DADA2 only to quality-check, filter, and trim sequences with the same parameters as for 16S gene but using a truncation length of 290 bp for the forward and 250 for the reverse complement. Using NifMAP, we merged reads in usearch v.9.2.64 (Edgar and Bateman 2010) and filtered sequences using hmmer v.3.3 (Eddy 2011) and a Hidden-Markov Model (*hmm_nuc_1160_nifH.hmm*). Sequences were dereplicated, and singletons were excluded from the analysis. In this case, we generated OTUs at 0.97% identity using usearch instead of ASVs. The sequences were translated to amino acids and corrected for frameshift errors using FrameBot v.1.2 (Wang et al. 2013). Then, the sequences were screened with hmmer (*nifH*_ChlL_bchX.hmm) to discard homologous genes, and a processed OTU table was generated. Finally, OTUs were classified using three approaches: the Evolutionary Placement Algorithm of RAxML (Stamatakis 2014), the Classification and Regression Trees (CART) to assign phylogenetic cluster of *nifH* genes (Frank et al. 2016) and using BLASTP (Camacho et al. 2009) against the RefSeq database (Pruitt et al. 2005). Library size and sample DNA concentrations were used to estimate potential contaminants in decontam package (Fig. S5). A 0.5% prevalence threshold was applied to filter out the resulting OTUs (Fig. S6).

### Bacterial community analyses

To compare *R. lanuginosum* bacterial composition associated with each tundra type, we transformed the 16S rRNA and *nifH* genes data taxa counts to relative abundances using phyloseq. The relative abundance per phylum and order was compared between the forest tundra and shrub tundra. We separated a subset of the 50 and 20 most abundant taxa for 16S rRNA and *nifH* genes, respectively, and compared their relative abundance per family between tundra types. We normalized the taxa table to compare alpha diversity between habitats using the relative abundance and multiplying it by a constant of 1000 to obtain count data. The Shannon, Simpson and Pielou indexes were estimated using the normalized tables. In addition, we calculated Faith’s phylogenetic diversity to consider the phylogenetic relationships of taxa using pairwise distances derived from a phylogenetic tree (Faith 1992). First, we aligned sequences in DECIPHER v.2.22.0 (Wright, Erik 2016) with default settings. Then, a phylogenetic tree was constructed using a neighbour-joining algorithm with a GTR+ GAMMA model in phangorn v.2.8.1(Schliep 2011) for both 16S and *nifH* data. Finally, we used the package picante v.1.8.2 (Kembel et al. 2010) to compute the phylogenetic diversity using the normalized data and the unrooted maximum-likelihood tree. We calculated the mean of each alpha diversity index per plot by averaging the values of the three squares or replicates. Shapiro normality tests were applied to choose the appropriate statistical test. We assessed differences in Shannon, Simpson, Pielou and Faith’s index of both datasets between habitats with the mean values per plot using Mann-Whitney tests in R v.4.1.3 (R Core Team 2017).

Beta diversity was assessed using Generalized Unifrac distances in the package GUniFrac v.1.4 (Chen et al. 2012). Generalized Unifrac has been proven effective when detecting community changes shaped by moderately abundant taxa (ASVs or OTUs) while identifying rare or abundant ones compared to the unweighted and weighted measures (Chen et al. 2012). The previously produced maximum-likelihood trees were used to construct the distance matrices with an alpha value of 0.5, as recommended in the package GUniFrac. The distance matrices were analyzed using a Non-Metric Multidimensional Scaling (NMDS) and Principal Coordinates Analyses (PCoA) in phyloseq. To test for statistical differences in microbial composition between habitats, we performed a permutational multivariate analysis of variance (PERMANOVA) with the distance matrices using *adonis* function and 9999 permutations in vegan v.2.5.7 (Oksanen et al. 2018). We added the plot as strata to account for the repeated measures. Finally, we verified the dispersion of groups to confirm the group heteroskedasticity assumption.

### Detection of bacteria associated with the moss, soil, and habitats

The core microbial community of *Racomitrium lanuginosum* was investigated using the R package microbiome v.1.16.0 (Lahti and Shetty 2017). We defined a core community as the ASVs and OTUs common to the moss species and tundra type using different prevalence thresholds and based on shared community composition (see Shade and Handelsman 2012; Alonso-García and Villarreal Aguilar 2022). The core bacteriome of the moss (16S rRNA and *nifH* genes) and soil were inferred using the command *core_members* with a minimum prevalence of 0.75 and a 0.01 detection threshold. We compared the moss and soil core microbiome to detect ASVs co-occurring in both samples. In addition, the moss core microbiome of the forest tundra and shrub tundra was investigated to identify potential bacteria associated with the two habitats. Results were plotted as Venn diagrams to visualize exclusive and shared core members with the package eulerr v.6.1.1 (Larsson 2022). Finally, we only focused on the moss core community and filtered ASVs/OTUs with a minimal prevalence of 0.75 and 0.90. The taxonomic information of the core members was recovered, and their prevalence was plotted as a response to the relative abundance detection threshold.

Bacterial and diazotrophic groups characteristic of each habitat were identified using linear discriminant analysis effect size using the Galaxymodule LEfSe (Segata et al. 2011). LEfSe uses the relative abundance of taxa as input and performs Kruskal-Wallis tests to identify differentially abundant taxa or groups (taxonomic ranks) between two conditions or treatments. If provided, differences among subclasses are estimated using Wilcoxon tests to investigate biological significance. Finally, linear discriminant analysis was applied to estimate the effect size of differentially abundant taxa (Segata et al. 2011). We formatted 16S rRNA and *nifH* genes abundance tables to LEfSe input files. We performed LEfSe tests at genus and ASV/OTU levels using only habitat type as classes (forest and shrub tundra) with default parameters (LDA threshold = 2.0). Differentially abundant groups and taxa are presented in cladograms and ranking plots for each dataset.

### Acetylene reduction assays

We employed acetylene reduction assay (ARA) (Hardy et al. 1968) to infer the N_2_-fixation of the moss-microbiome. Three moss replicates per plot were placed in 23 ml glass vials, humidified, and acclimated at room temperature (∼21°C) for 24h before experiments. We cut the brown tissue excess in the basal section of the shoots. Acetylene gas was produced by combining 25 ml of H_2_O and 5 g of CaC_2_ in a gas sampling bag with a septum. From each vial, 3 ml of air was replaced with acetylene for a final concentration of ∼20%. Samples were incubated for 24h at 23°C in continuous light. After incubation, we collected 3 ml of gas from the vials to measure ethylene production using a GC-FID (Shimadzu 8A) equipped with a Supelco column 01282011 (Supelco Analytical) and using helium as carrier gas. To transform the ethylene area into concentrations in ppm, we utilized calibration curves of ethylene at 2.5, 5, 10, 50 and 100 ppm. Then, moss samples were oven-dried at 50°C for 24 h to obtain the dry weight. Acetylene reduction rates are reported in nmol C_2_H_4_/h/g.

### Moss element analyses

To explore the relationship between the moss-microbiome N_2_-fixation rates and nutrient content in moss, we performed element analyses of the same dried samples used for the ARAs. We followed the moss element concentration analyses described in Renaudin et al. (2022a). Dried samples were manually ground to a fine powder using a mortar and pestle in liquid nitrogen. Around 50 mg of each sample was put in trace metal-free tubes (SCP Sciences) with 1 ml of nitric acid (trace metal-free grade, ThermoFisher Scientific) and 200 μL of hydrogen peroxide (trace metal-free grade, MilliporeSigma). Digestion was performed at room temperature for 30 minutes, followed by 1 h at 45°C and 2 h at 65°C in a heating block digestion system (DigiPREP, SCP Sciences). After digestion, an acid concentration of 2% was reached by adding Mili-Q water (MilliporeSigma). Element concentrations were measured on an ICP-MS (X-Series II, ThermoFisher Scientific) using rhodium (Rh) as the internal standard. Element concentrations were transformed to ppm using the dried weight of the samples. We focused on the nitrogenase-related nutrients P, Fe, V and Mo.

### Statistical analyses

Comparisons of the acetylene reduction rates and nutrient content between tundra types were performed by averaging values per plot, as in alpha diversity analyses, and using them as input. In addition, Gaussian distribution of data was assessed with Shapiro tests. We conducted Mann-Whitney tests between habitats for all variable comparisons in the R environment (acetylene reduction rates and moss nutrient content).

The response of the moss-associated N_2_-fixation rates to the relative abundance of the 20 most abundant diazotrophic genera (*nifH* gene) was analyzed. The log-normal distribution was selected *a priori* by fitting acetylene reduction rates against different theoretical distributions. The glmmTMB package v.1.1.3 (Brooks, Mollie et al. 2017) based on maximum likelihood was used to analyze the data. We used the mean values of N_2_-fixation rates and diazotrophic abundance per plot to fit the model. Then, Generalized Linear Models (GLM) were carried out using the moss-microbiome N_2_-fixation rates as a response and the relative abundance of the following dominant diazotrophic genera as predictors: *Anabaena*, *Azorhizobium*, *Calothrix*, *Dolichospermum*, *Fischerella*, *Iningainema*, *Mastigocladus*, *Nostoc*, *Rhodomicrobium* and *Trichormus*. The model was constructed on glmmTMB with a normal error distribution and a log link function.

We also investigated the relationships of acetylene reduction rates (N_2_-fixation) with the moss nutrient content and habitat. We removed data outliers and scaled moss nutrient content (*scale* function) to have comparable values. N_2_-fixation and moss nutrient content were measured with three replicates per plot. Hence, all replicates were used to construct models by controlling for variation within plots. We applied Generalized Linear Mixed-effects Models (GLMM) using glmmTMB. N_2_-fixation rates were treated as the response variable, moss nutrient content (P, Fe, V, Mo) and habitat type as fixed factors, and plots as a random factor with a normal error distribution and a log link function.

All models were validated by examining the residuals’ homogeneity and normal distribution of errors. In addition, for GLMMs, we inspected the random effects distribution. All statistical analyses were carried out in R.

## RESULTS

### Bacterial diversity and community structure

The filtered 16S rRNA data of *Racomitrium lanuginosum* and associated soil had a mean library size of 9387 reads per sample, resulting in 4885 ASVs from 89 samples (62 for the moss and 27 for the soil). The microbial community of *R. lanuginosum* was represented by 1322 ASVs and mainly dominated by the phyla Proteobacteria (28.20%), WPS-2 or Eremiobacterota (19.21%), Actinobacteria (12.94%), Chloroflexi (12.83%), Acidobacteria (9.56%) and Cyanobacteria (7.57%) (Fig. 2a; Fig. S7). The more abundant bacterial orders were Acetobacterales, Acidobacterales, Isosphaerales, Ktedonobacterales and Frankiales (Fig. S8). The 50 most abundant ASVs grouped bacterial families such as Acetobacteraceae, an unclassified group of Eremiobacterota, Ktedonobacteraceae, Nostocaceae and Isosphaeraceae (Fig. S9). Soil bacterial community was primarily composed of Proteobacteria (26.11%), Actinobacteria (21.78%), Acidobacteria (17.90%), Chloroflexi (11.96%), Planctomycetes (7.62%) and Eremiobacterota (4.40%) (Figs. S10 and S11). The 50 most abundant ASVs included Acidothermaceae, Ktedonobacteraceae, Solibacteraceae (Subgroup 3) and unclassified members of Frankiales (Fig. S12).

**Fig. 2.**
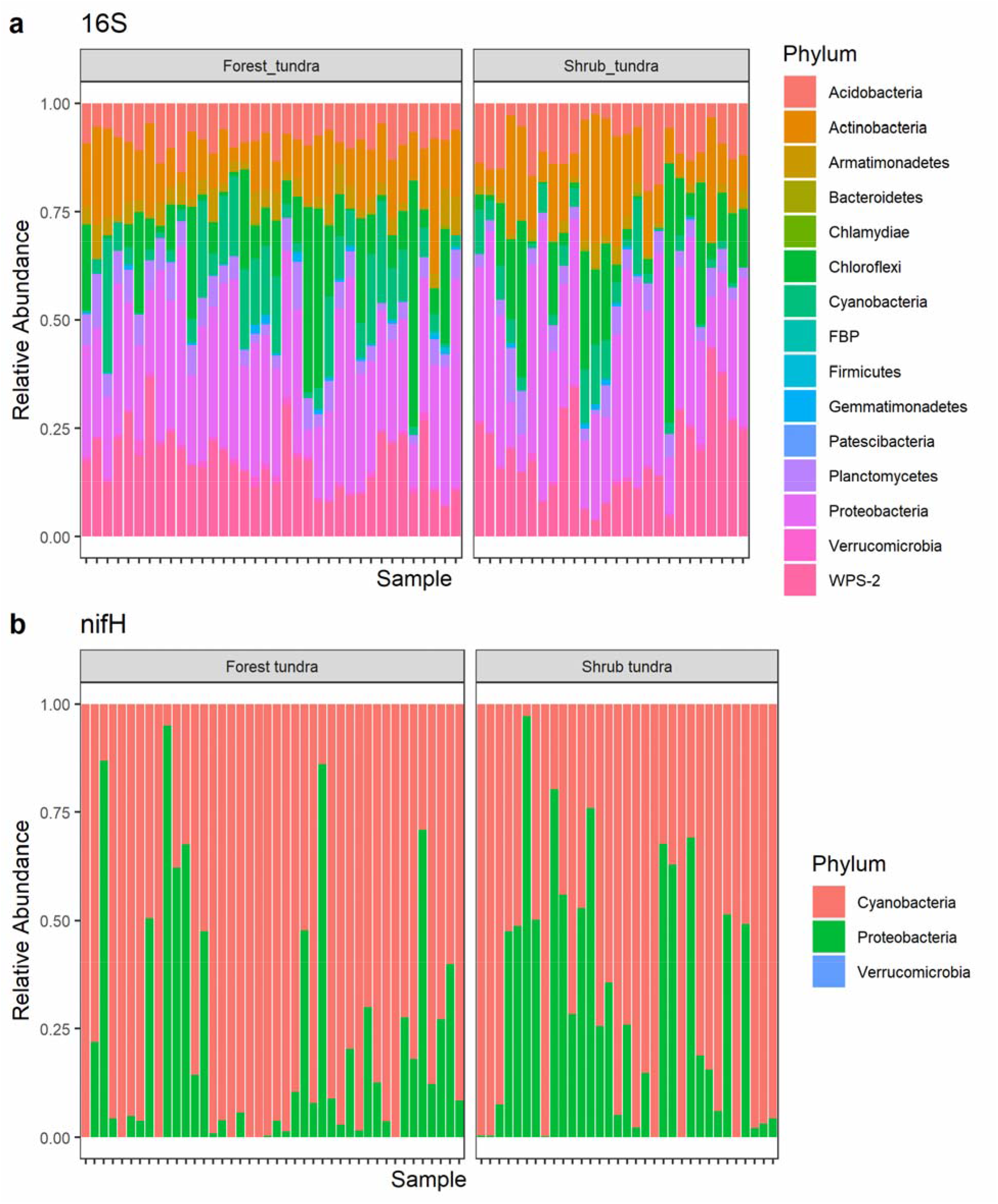
Relative abundance of *Racomitrium lanuginosum* associated bacterial phyla based on 16S rRNA and *nifH* genes. The relative abundance is presented as proportions on the x-axis and samples on the y-axis, each bar represents one sample. Colours indicate each phylum in the legend. **a** Relative abundance based on 16S rRNA gene in two tundra types. The most abundant phyla are Proteobacteria, Actinobacteria, Acidobacteria and Planctomycetes. Samples are divided into forest tundra (Kuujjuarapik, n=36) and shrub tundra (Umiujaq; n=26). **b** Relative abundance based on *nifH* gene in two tundra types. The most abundant phyla are Cyanobacteria and Proteobacteria. Samples are divided into forest tundra (Kuujjuarapik; n= 42) and shrub tundra (Umiujaq; n= 33).

Amplicon data of the *nifH* gene showed a mean library size of 5509 reads with 105 processed OTUs in 75 samples. The diazotrophic diversity hosted by *R. lanuginosum* was dominated by Cyanobacteria (67.86%) and Proteobacteria (32.13%), with a predominance of the family Nostocaceae (Fig. 2b; Fig. S13). The 20 most abundant OTUs inhabiting the moss included the Nostocaceae and Aphanizomenonaceae (Fig. S14). In addition, we found that diazotrophs belonging to *nifH* phylogenetic clusters 1B and 1K were abundant. These groups include taxa having the Mo-nitrogenase, comprising Cyanobacteria (1B) and members of Alphaproteobacteria (1K), such as *Azorhizobium* and *Rhodomicrobium* (Fig. S15).

Alpha diversity estimates suggested that moss bacterial communities are composed of a few dominant taxa and many others with low abundance. The soil bacterial communities based on 16S rRNA gene were the most diverse, followed by the *R. lanuginosum* bacterial (16S rRNA) and diazotrophic (*nifH*) communities (Fig. 3). According to Mann-Whitney tests, Shannon and Simpson’s indexes did not differ between tundra types. However, Faith’s phylogenetic diversity of the moss microbiome (U = 115, p-value = 0.007) and Pielou evenness of the soil bacterial community (U=136, p-value=0.024) were higher in the forest tundra than in the shrub tundra.

**Fig. 3.**
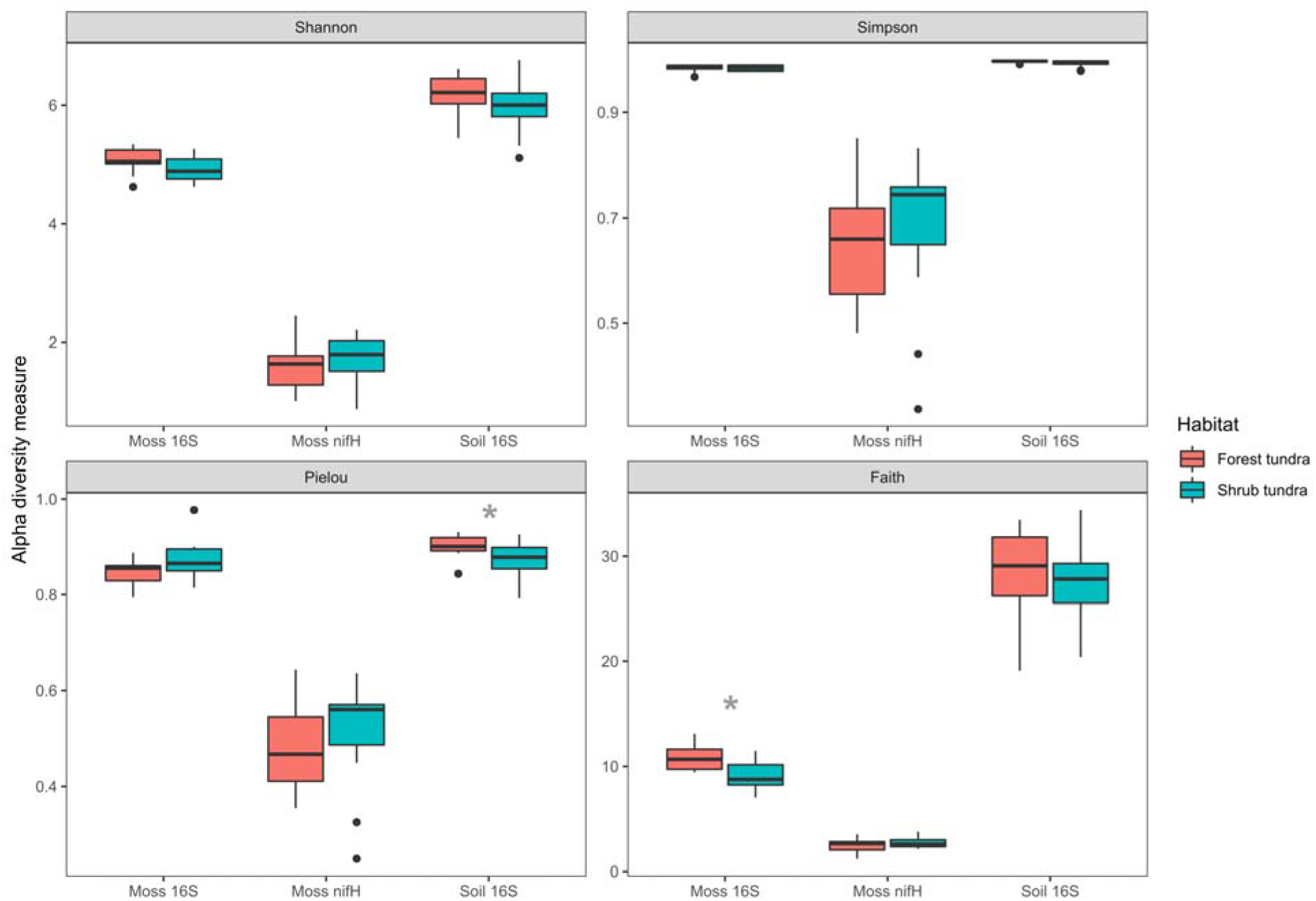
Alpha diversity indexes based on bacterial taxa (16S rRNA and *nifH* genes) associated to the moss *Racomitrium lanuginosum* and associated soil in two types of tundra. Shannon, Simpson, Pielou and Faith’s phylogenetic indexes were estimated by averaging values per plot in the case of moss data. The box represents the first quartile, median and third quartile, with whiskers indicating the maximum and minimum values and outliers presented as points for each dataset in two habitats. Grey asterisks indicate significant differences between habitats (p-value < 0.05). Only moss 16S Faith’s phylogenetic diversity and Pielou’s evenness of the soil bacteria are higher in the forest tundra

Beta diversity analyses of ASVs/OTUs using PERMANOVAs indicated no significant differences between *R. lanuginosum* bacterial and diazotrophic community composition between tundra types when controlling for plot variation (Table S2). Accordingly, the tundra type only explained 4% of moss bacterial and diazotrophic assemblages’ variation. The ordination analyses did not reveal any apparent tundra-type clustering for both moss datasets; however, the bacterial composition of the soil was affected by the habitat type (Figs. S16-S18).

LEfSe tests inferred bacterial and diazotrophic taxa explaining the differences between the two tundra types. The bacterial genera related to the forest tundra comprised several groups, such as *Nostoc*, *Bryocella*, *Kinesporia*, *Tepidisphaera*, *Fimbriiglobus* and *Methylobacterium*. In contrast, the shrub tundra was mainly characterized by the order Acetobacterales (*Acidiphilium* and *Acidocella*) and the genus *Granulicella* (Fig. 4; Fig. S19). Regarding the diazotrophic community, only the genera *Calothrix* and *Anabaena* were differentially abundant in the shrub tundra (Fig. 4; Fig. S19). We also analyzed the diazotrophic community at the OTU scale and found that each habitat was characterized by potential diazotrophs belonging to similar taxonomic groups as in the genus-scale analysis (Fig. S20).

**Fig. 4.**
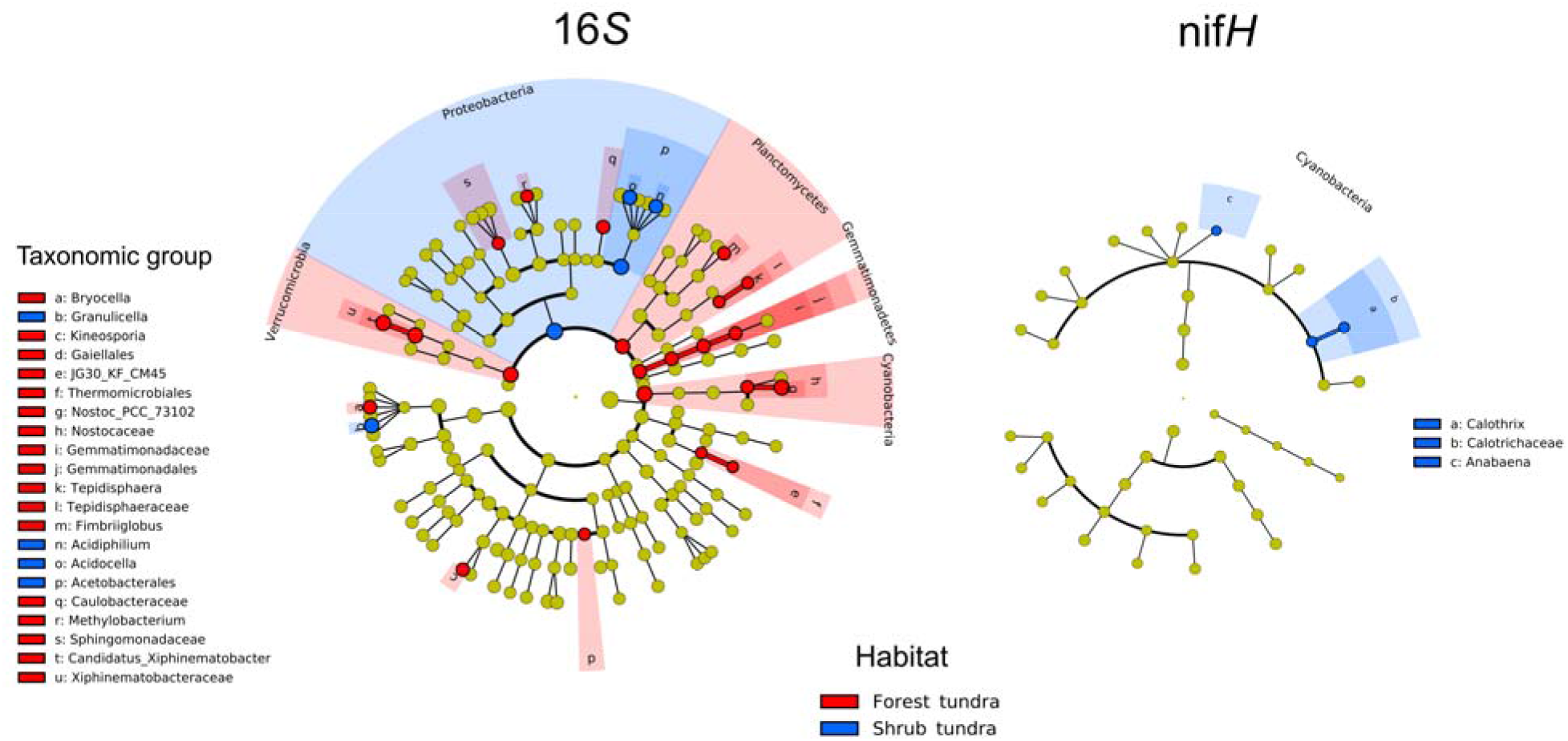
Cladograms of the differentially abundant bacterial genera associated to *Racomitrium lanuginosum* in two habitats based on LEfSe analyses. On the left, a cladogram of the bacterial community based on 16S rRNA gene shows the differentially abundant groups for the forest tundra in red and the shrub tundra in blue. The legends indicate taxonomic groups with letters within the cladograms. The forest tundra is characterized by several genera, while Proteobacteria mainly denote the shrub tundra. On the right, a cladogram for the diazotrophic community based on the *nifH* gene shows the differentially abundant groups using the same color code for each habitat. *Anabaena* and *Calothrix* were the only diazotrophic genera distinctly abundant in the shrub tundra

### Core microbiome

The core microbiome of *R. lanuginosum* based on the 16S rRNA gene dataset is composed of 29 ASVs (prevalence threshold >0.75), and no core bacteria were shared with its surrounding soil (Fig. 5a). The moss core bacterial community in both habitats was similar. Still, five and four ASVs were more related to the forest and shrub tundra, respectively. The *R. lanuginosum* core bacteria were represented by *Acidiphilium*, *Singulisphaera*, *Bryobacter* and unclassified taxa from WPS-2 (Eremiobacterota) and Acidobacteraceae. The moss core community in the forest tundra comprised ASVs of *Acidiphilum*, two unidentified ASVs from Acetobacteraceae, *Conexibacter* and *Granulicella*; while in the shrub tundra, the core bacteria were *Conexibacter*, *Acidisphaera* and unnamed ASVs in Beijerinckiaceae and Eremiobacterota. As expected, core microbiome analysis using a more stringent prevalence threshold (>0.90) reduced the core microbiome. Here, only 14 ASVs belonged to the same groups (Fig. S21).

**Fig. 5.**
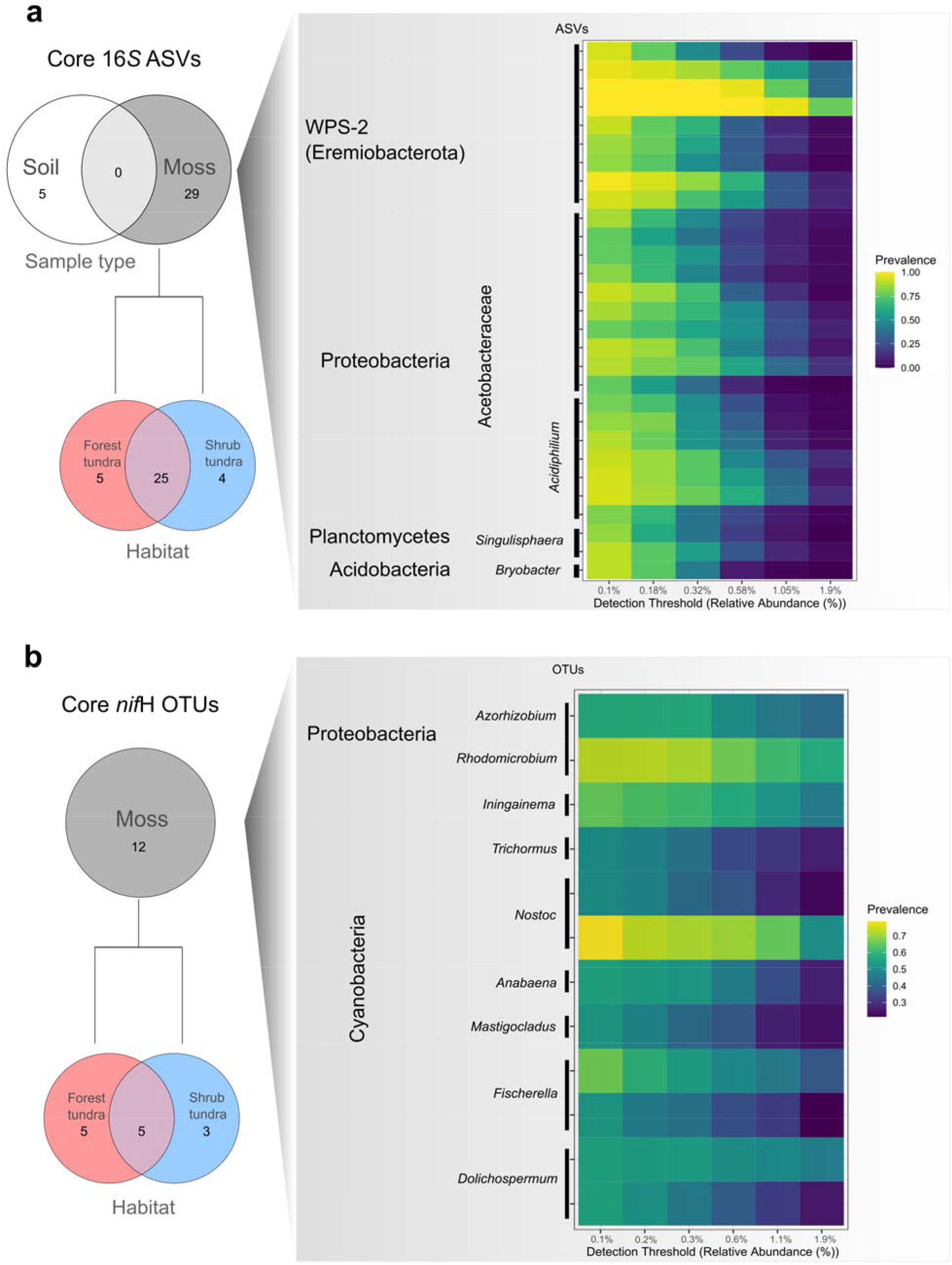
Core microbiome of *Racomitrium lanuginosum*. **a** The left panel shows Venn diagrams based on 16S rRNA gene data with the number of core ASVs for the moss *R. lanuginosum* and its associated soil and moss core ASVs for each habitat. The right panel includes the *R. lanuginosum* core microbial community and a heatmap with the ASV prevalence in response to the relative abundance detection threshold. Core ASVs are presented on the y-axis with their available taxonomic information and a minimum prevalence detection threshold of 0.75. The color scale on the left indicates the ASV prevalence from minimum to higher values on a blue-to-yellow scale. Increasing the detection threshold will reduce the prevalence and, thus, the number of core ASVs. The candidate phylum WPS-2 (Eremiobacterota) is a common group of the *R. lanuginosum* microbiome. **b** Core microbiome based on *nifH* gene: A Venn diagram with the moss core diazotrophic OTUs in two tundra sites. On the right panel, the core diazotrophic bacteria and their taxonomic information is presented with a minimum detection threshold of 0.50. Note that the detection threshold is lower for the diazotrophic community, but the color scale is represented similarly to the 16S rRNA gene data. Cyanobacteria dominate the diazotrophic core community

The analysis of the core diazotrophic bacteria of *R. lanuginosum* with a 0.75 prevalence threshold recovered only one *Nostoc* OTU; However, more diazotrophic genera were retrieved when attenuating the threshold to 0.50 (Fig. 5b). The diazotrophic core community at 0.50 prevalence threshold included members of Cyanobacteria such as *Anabaena*, *Fischerella*, *Dolichospermum* and *Mastigocladus* and OTUs from Proteobacteria such as *Azorhizobium* and *Rhodomicrobium*. We found slightly different core diazotroph genera in each tundra type. The core diazotrophic bacteria for the forest tundra were *Dolichospermum*, *Trichormus*, *Mastigocladus*, *Fischerella* and *Nostoc*, while the shrub tundra has a diazotrophic community represented by *Anabaena*, *Nostoc* and *Dolichospermum* OTUs.

### N_2_-fixation rates and associated variables

The estimated mean acetylene reduction rate of *R. lanuginosum* was 21.86 nmol C_2_H_4_/h^-1^/g^-1^. Acetylene reduction rates varied according to the tundra type, with the forest tundra having a mean rate of 16.75 C_2_H_4_/h^-1^/g moss^-1^ (±15.77) and the shrub tundra 27.54 C_2_H_4_/h^-1^/g moss^-1^ (±20.85) (Fig. 6). Nonetheless, we did not detect statistically significant differences between habitats. The relative abundances of some of the 20 most dominant diazotrophic genera (Fig. S14), such as *Azorhizobium*, *Calothrix*, *Dolichospermum*, *Fischerella*, *Nostoc* and *Rhodomicrobium,* were significantly related to the N_2_-fixation rates (Table 1). These results suggest an essential role of diazotrophs in driving the N_2_-fixation of *R. lanuginosum* in the tundra. Interestingly, *Azorhizobium* and *Rhodomicrobium* are reported for the first time to be statistically associated with N_2_-fixation rates in mosses and may represent essential contributors to the tundra N inputs.

**Fig. 6.**
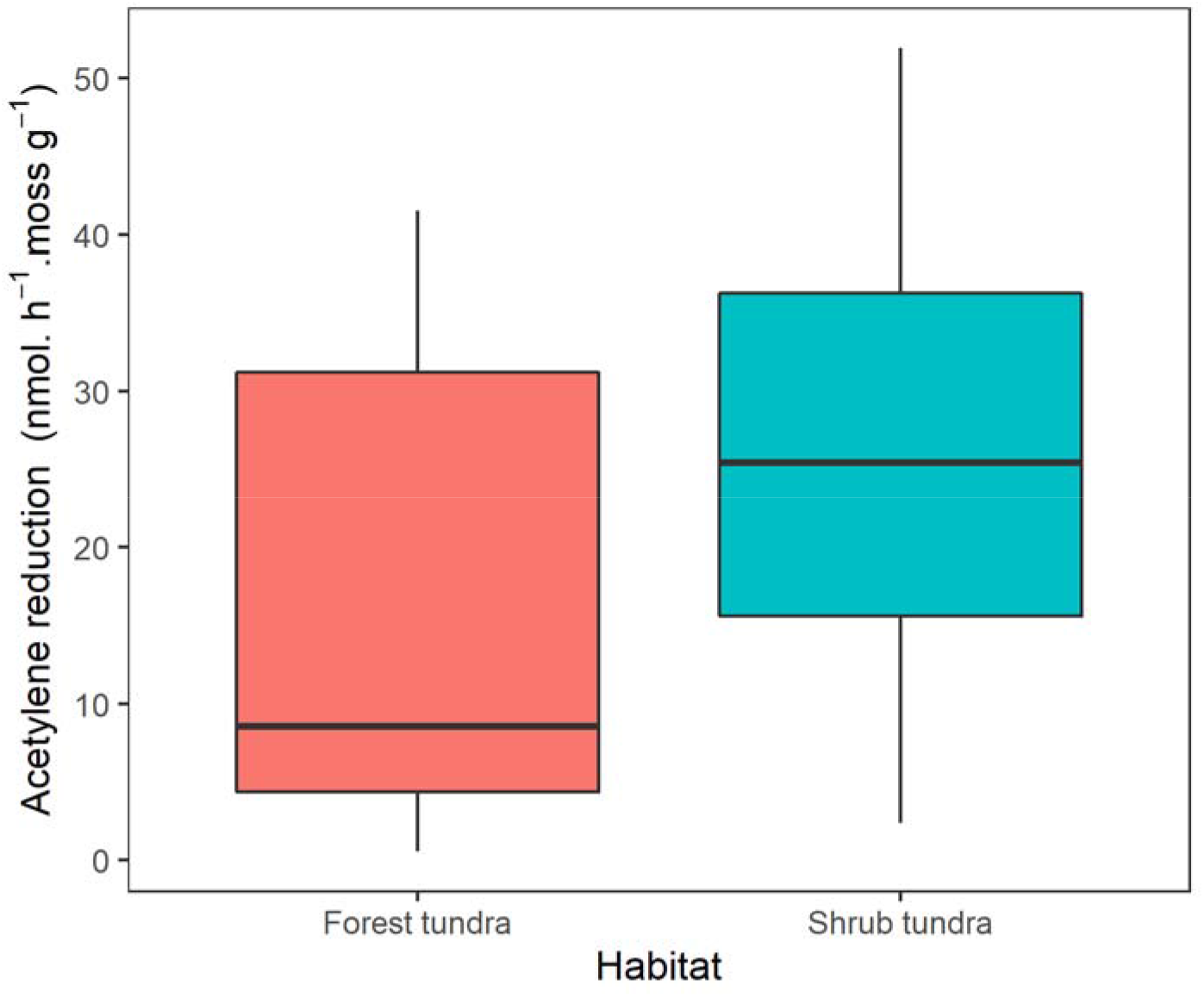
Acetylene reduction rates associated with *Racomitrium lanuginosum* in two types of tundra. The boxplot represents the first quartile, median and third quartile, with whiskers indicating the maximum and minimum values (n=13). Acetylene reduction rates (N_2_-fixation) did not significantly differ between tundra types

**Table 1.** Generalized linear model of *Racomitrium lanuginosum*-microbiome acetylene reduction rates as a response to the abundance of the dominant diazotrophic bacterial genera and “log” as link function. The estimates, standard error (SE), z value and p-value are presented. The model was fitted using 93 observations. The abundance of the diazotrophic genera *Azorhizobium*, *Calothrix*, *Dolichospermum*, *Fischerella*, *Nostoc*, and *Rhodomicrobium* have a significant statistical effect on moss-associated N_2_-fixation rates

|  | Estimate | SE | z | p-value |
| --- | --- | --- | --- | --- |
| (Intercept) | -0.8076 | 1.9061 | -0.424 | 0.6718 |
| *Anabaena* | -27.7563 | 17.5964 | -1.577 | 0.1147 |
| *Azorhizobium* | 6.3982 | 2.3811 | 2.687 | 0.0072 ** |
| *Calothrix* | 5.6049 | 2.4992 | 2.243 | 0.0249 * |
| *Dolichospermum* | 4.3462 | 1.961 | 2.216 | 0.0266* |
| *Fischerella* | 11.4603 | 5.418 | 2.115 | 0.0344* |
| *Iningainema* | -4.5217 | 6.8161 | -0.663 | 0.5071 |
| *Mastigocladus* | -11.6351 | 7.4356 | -1.565 | 0.1176 |
| *Nostoc* | 5.3325 | 2.5651 | 2.079 | 0.0376 * |
| *Rhodomicrobium* | 5.3372 | 2.5553 | 2.089 | 0.0367 * |
| *Trichormus* | 2.6104 | 2.1755 | 1.2 | 0.2302 |

\* $p$ -value $< 0.05$ ; \*\* $p$ -value $< 0.01$ .

The nutrient concentrations in *R. lanuginosum* were similar in both tundra types with no statistical differences (Fig. 7). The regression models of moss nutrient content on moss-associated acetylene reduction rates indicated that P, Fe, V, and Mo do not statistically affect the ability of the *R. lanuginosum* microbiome to fix nitrogen (Table 2). Thus, we inferred that nutrients might have little effect on *R. lanuginosum*’s BNF in the forest-tundra transition zone. In addition, GLMM confirmed similar N_2_-fixation rates between habitats.

**Fig. 7.**
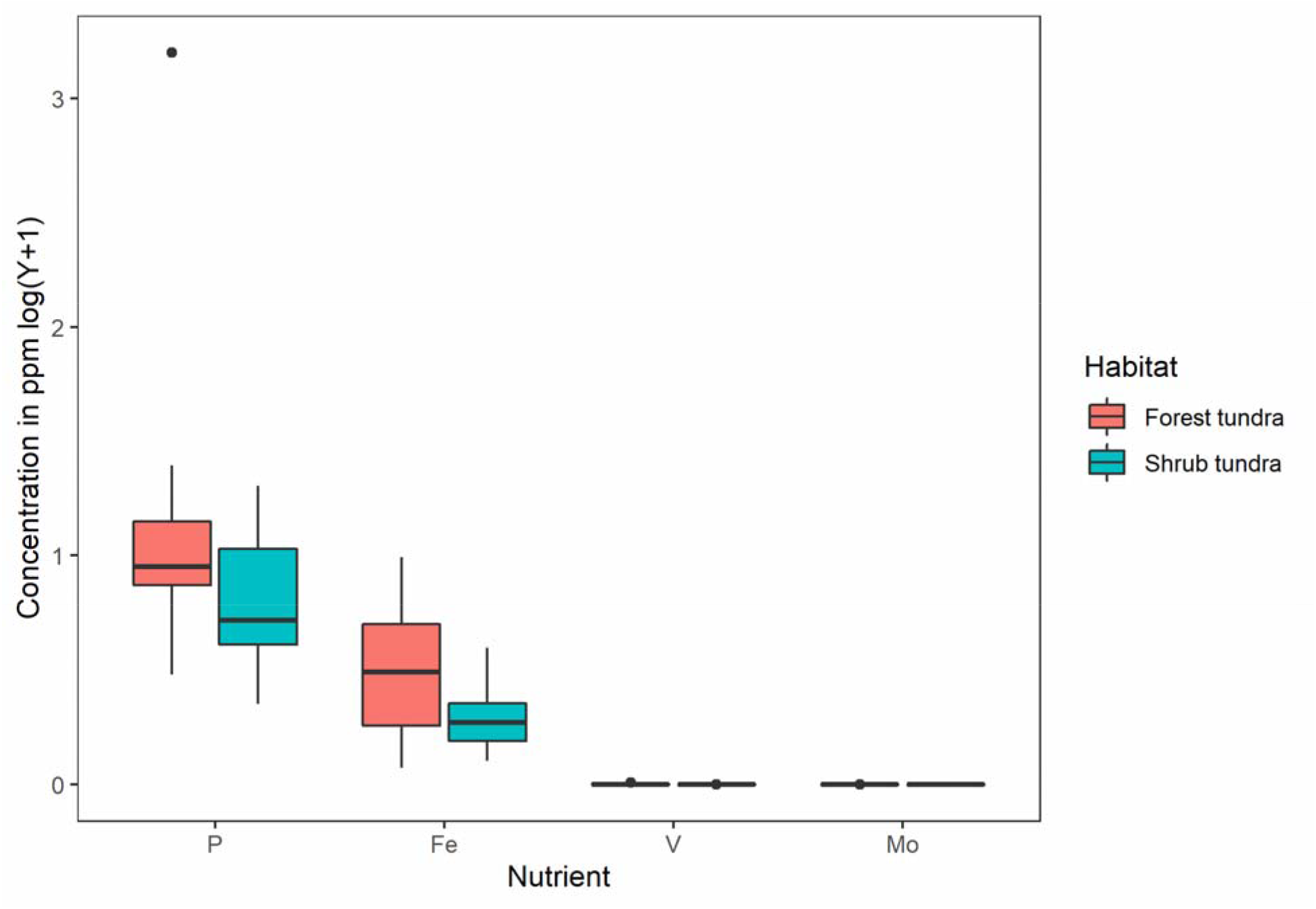
Nutrient content of *Racomitrium lanuginosum* in two types of tundra. The box represents the first quartile, median and third quartile, with whiskers indicating the maximum and minimum values and outliers presented as points (n=12). No statistical differences were found between moss nutrient concentrations in the two habitats

**Table 2.** Generalized linear mixed-effect model of *Racomitrium lanuginosum*-microbiome acetylene reduction rates as a response to nitrogenase-related nutrient content of the moss (P, Fe, V, Mo) and habitat type (fixed effects) with plots included as a random effect and “log” as link function. The estimates, standard error (SE), z value and p-value are presented. The model was fitted using 64 observations grouped in 24 plots. The standard deviation of plots (SD Plot) represents the variation of random effects. The moss nutrient content and tundra type do not have a significant statistical effect on moss-associated N_2_-fixation rates

|  | Estimate | SE | z | p-value |
| --- | --- | --- | --- | --- |
| Intercept | 2.4462 | 0.2720 | 8.994 | $< 2e-16$ *** |
| SD Plot | 0.7533 |  |  |  |
| P | -0.1904 | 0.1528 | -1.246 | 0.213 |
| Fe | 0.1191 | 0.1676 | 0.711 | 0.477 |
| V | -0.0524 | 0.1946 | -0.269 | 0.788 |
| Mo | 0.0279 | 0.0855 | 0.326 | 0.744 |
| Habitat-Shrub Tundra | 0.5286 | 0.3529 | 1.498 | 0.134 |
| *** p-value <0.001= |  |  |  |  |

## DISCUSSION

In this research, we characterized the microbial community of the moss *Racomitrium lanuginosum* in the forest-tundra transition zone. We found differential bacterial groups that are abundant in each tundra type. The species core microbiome was characterized by bacteria with potential roles in moss symbiosis. Finally, we report new groups of diazotrophs significantly correlated with moss-bacteria N_2_-fixation rates in this cold environment.

### Similar diversity but differentially abundant bacteria between tundra types

In contrast to the initial hypothesis that the microbial community in *R*. *lanuginosum* responds to habitat type, we found that most of the bacterial members in microbial communities of the forest tundra and shrub tundra were similar. However, the abundance of certain bacterial groups differed between habitats. The core microbiome analyses per habitat and the LDA results indicated abundant differential genera in the forest tundra and shrub tundra.

The forest tundra was characterized by the cyanobacterium *Nostoc* PCC-73102 (Fig. S19). This bacterium was also significantly abundant in feather mosses, and its abundance positively correlates with tree density in boreal forests of Eastern Quebec (Renaudin et al. 2022b). The prevalence of Nostocaceae in the forest tundra also might relate to its temperature optimum for N_2_-fixation, which is reported to be around 13°C (Gentili et al. 2005). In contrast, the high abundance of some Proteobacteria in *R. lanuginosum* in the shrub tundra may be influenced by the dwarf shrub biomass and labile litter, as reported for Icelandic populations (Klarenberg et al. 2021). Other diazotrophs, such as *Calothrix* and *Anabaena,* were identified as differentially abundant in the shrub tundra. Both genera are common cyanobacterial symbionts in bryophytes and some vascular plants (Warshan et al. 2016, 2017; de Jesús Suárez-Moo et al. 2019; Bouchard et al. 2020; Holland-Moritz et al. 2021; Alvarenga and Rousk 2022; Renaudin et al. 2022b).

Environmental conditions allow some bacteria associated with *R. lanuginosum* to become abundant in each tundra type. A temporal record of microbial diversity is desired to identify the abundance and bacterial composition differences between habitats accurately.

### The core microbiome is associated with northern mosses with limited soil bacterial recruitment

Contrary to our hypothesis, the *R. lanuginosum* core microbiome did not share bacteria from its soil. We stressed that soil bacterial recruitment to the moss indeed occurs but not at the level of the core microbial community (prevalence >0.75). For example, the basal part of feather moss shoots is composed of bacteria known from forest soils and with degradation ability, while the apical portion (photosynthetic) is dominated by cyanobacteria which have better access to N_2_ and light (Renaudin et al. 2022b).

Instead, the core bacteria members may come from other environmental sources (water currents, air) or are most likely to be shared with sympatric species (mosses, vascular plants or lichens), as exemplified in the microbial communities of lichens and trees from northern ecosystems (Lajoie and Kembel 2021; Alonso-García and Villarreal Aguilar 2022). For instance, the core bacterial community of *R. lanuginosum* is formed of common moss-symbiotic bacteria, such as Proteobacteria, Actinobacteria, Acidobacteria, Eremiobacterota and Cyanobacteria (Kostka et al. 2016; Warshan et al. 2016; Holland-Moritz et al. 2018, 2021; Renaudin et al. 2022a, b; RodríguezLJRodríguez et al. 2022). The strong association of these bacteria with other moss species may suggest a widespread microbial consortium that performs diverse functions in symbiosis with northern mosses. For example, among the *R. lanuginosum* core bacteria, the genus *Bryobacter* (Acidobacteria) is an aerobic chemo-organotroph strongly related to *Sphagnum* microbial communities (Kulichevskaya et al. 2010). This bacterium is involved in the C cycling and may provide nutrition to the moss. Similarly, the genus *Singulisphaera* (Planctomycetes) is also associated with *Sphagnum* spp. and is a chemo-organotrophic aerobe with a potential ability for secondary metabolite synthesis (Kulichevskaya et al. 2008; Morrow et al. 2020). This protobacterial genus *Acidiphilium* has been reported from mining-related water and is characterized by reducing ferric iron under anoxic and microaerobic conditions and maintains an association with *Acidithiobacillus ferrooxidans,* where the first transfer carbon dioxide to the latter (Ullrich et al. 2015). The most abundant group of the *R. lanuginosum* core microbiome is the recently proposed phylum WPS-2 (Eremiobacterota), widely distributed in cool, acidic, and aerobic environments, such as bogs, and with the ability to perform anoxygenic photosynthesis (Ward et al. 2019). These bacteria have been reported to be abundant in symbiosis with boreal mosses and reindeer lichens and they could be essential components of nutrient cycling in subarctic settings (Holland-Moritz et al. 2018; Alonso-García and Villarreal Aguilar 2022). The congruence of the moss core microbiome is kept when comparing the microbiome of *R. lanuginosum* of the forest-tundra transition zone in Eastern Canada to that of Icelandic populations sharing abundant bacterial taxa: *Bryobacter*, *Acidiphilium*, *Granulicella*, *Conexibacter* and *Nostoc* (Klarenberg et al. 2021). These similarities suggest a relatively stable species core microbiome across populations, even on large geographic scales.

### Abundant diazotrophs drive N_2_-fixation at the local scale

The acetylene reduction rates between the forest and shrub tundra moss populations did not differ statistically in this study. Similar results were found for the N_2_-fixation rates of *H. splendens* contrasting populations in the boreal forest and arctic tundra (Liu and Rousk 2022) and for foliar endophytic bacteria of conifers of the forest tree line and a subalpine forest (Moyes et al. 2016). The bacterial community composition of *R. lanuginosum* is similar in the forest tundra and shrub tundra (Figs. S16 and S17), which results in similar N_2_-fixation rates.

The mean acetylene reduction rate of *R. lanuginosum* in the forest-tundra transition zone was 21.86 nmol C2H4/h-1/g moss-1. Even though there are previous reports of the species N_2_-fixation activity, direct comparison is not feasible due to different scales. If we transform our data into N_2_ using a conversion factor of 3 as reported in DeLuca et al. (2002), we obtain a rate of 7.28 nmol N/ h^-1^/g moss^-1^. For *R. lanuginosum* populations in the Alaskan arctic tundra, a rate of 11.69 ± 3.83 µg N/g moss^-1^/day^-1^ has been reported (34.75 ± 11.37 nmol N/ h^-1^/g moss^-1^) (Stuart et al. 2021). The N_2_-fixation rates per area of the congeneric *R. elongatum* Ehrh. ex Frisvoll in Washington, USA, are around 73.50 ± 31.74 µmol C_2_H_4_/m^-2^/day^-1^ (Calabria et al. 2020). The estimated *R. lanuginosum* N_2_-fixation rates seem lower than those reported in boreal feather mosses (Renaudin et al. 2022a); however, this moss is a conspicuous plant in the tundra and covers large areas of bare rock compared to other species, representing an essential contribution to the subarctic and arctic N budget.

The hypothesis that the abundant diazotrophic genera are essential predictors of N_2_-fixation rates holds true for *R. lanuginosum* in the forest-tundra ecotone. The bacterial composition at the shoot level plays an essential role in regulating N_2_-fixation. Among the identified bacterial groups, Nostocaceae is well known to associate positively with N_2_-fixation in mosses, but other bacteria from Verrucomicrobia, Planctomycetes, Actinobacteria, and Proteobacteria have been recently revealed (Jean et al. 2020; Holland-Moritz et al. 2021). In addition, the diazotrophic bacteria strongly correlated to N_2_-fixation in *R. lanuginosum* mainly include genera of Nostocales such as *Fischerella*, *Nostoc*, *Calothrix* and *Dolichospermum,* and the protobacterial genera *Azorhizobium* and *Rhodomicrobium*. In contrast, in boreal feather mosses (*P. schreberi*, *P. crista-castrensis* and *H. splendens*), cyanobacteria such as *Nostoc*, *Hassallia, Stigonema*, *Scytonema* and *Nodullaria,* and the proteobacterial taxa *Methylocapsa* and *Methyloferula* influence acetylene reduction rates (Leppänen et al. 2013; Renaudin et al. 2022b). These differences suggest alternative N_2_-fixing bacteria contributing to the N cycle in the forest-tundra zone.

Interestingly, the genera *Azorhizobium* and *Rhodomicrobium* have not yet been recognized as critical to N_2_-fixation in moss-bacterial symbioses in subarctic ecosystems. The genus *Azorhizobium* can be found as free-living or as a symbiotic bacterium of legume nodules, and it is used in agriculture; for example, the species *A. caulinodans* is an N_2_-fixer, hydrogen-oxidizer and common symbiont of a tropical legume (*Sesbania rostrata* Brem.) that forms stem nodules (Lee et al. 2008). Similarly, *Rhodomicrobium* spp. are freshwater bacteria with anoxygenic photoheterotrophic and photoautotrophic metabolisms (Ainon et al. 2006). N_2_-fixation tests indicate that *R. vannielii* can fix N_2_, even if the rates are lower than, for example, some *Rhodopseudomonas* strains (Madigan et al. 1984). Therefore, we postulated that in addition to Cyanobacteria, *Azorhizobium* and *Rhodomicrobium* play a part in fixing N_2_ under anoxygenic and light-limited conditions in *R. lanuginosum* populations of the forest-tundra zone.

Finally, the moss nutrient content does not significantly affect the N_2_-fixation rates, contrary to our hypothesis. These findings differed from studies on a large geographical scale, where nitrogenase-related elements such as P, Mo, and V correlate with N_2_-fixation rates in mosses and lichens (Rousk et al. 2017a; Darnajoux et al. 2019; Rousk and Rousk 2020; Rousk 2022; Renaudin et al. 2022a). Particularly, the effect of moss Mo, V and P content on N_2_-fixation is significant across an element deposition gradient in Eastern Canada (Renaudin et al. 2022a). For the forest-tundra zone, similar moss nutrient concentrations might have concealed any significant effect on N_2_-fixation rates. Thus, when nutrient concentrations are homogeneous or limited in the environment, the abundance of diazotrophs better predicts the local N_2_-fixation of the moss-bacteria symbiosis.

## CONCLUSIONS

The findings of this research reveal that the *Racomitrium lanuginosum* microbial community did not strictly respond to habitat type, with the forest tundra and shrub tundra having similar bacterial diversity but differentially abundant groups. Furthermore, the core moss microbiome does not share bacteria from its substrate despite the high soil microbial diversity. The *R. lanuginosum* core microbiome suggests a moss-associated bacterial consortium. Notably, *R. lanuginosum* has a recurrent association with uncharacterized strains of the recently proposed bacterial group WPS-2 (Eremiobacterota). Finally, we found that the moss-associated N_2_-fixation rates are similar between tundra types, but they seem to be performed by habitat-specific abundant bacteria. The abundance of some diazotrophs, not moss nutrient content, significantly affects *R. lanuginosum* N_2_-fixation locally in the forest-tundra transition zone. Notably, the proteobacterial genera *Rhodomicrobium* and *Azorhizobium* contribute to moss N_2_-fixation rates in the tundra. We recommend that future research could focus on experimentally assessing bacterial community changes facing the advance of the tree line into arctic environments. Overall, we identified critical bacterial groups related to moss-bacterial symbiosis and N_2_-fixation in the forest-tundra transition zone, a changing environment susceptible to climate warming.

## ACKNOWLEDGEMENTS

We thank the Center of Northern Studies (CEN) for the facilities provided in Kuujuarapik and Umiujaq. In addition, we would like to thank the Bellenger Lab for all the help with N_2_-fixation experiments, especially Ayan Ibrahim, for her valuable help with the moss element analyses.

## STATEMENTS AND DECLARATIONS

### Author contributions

The study and sampling design was conceived by JCVA, ND and DAEO. Sampling, molecular lab work and data analyses were performed DAEO. Acetylene reduction assays and element analyses were carried out by CB, JPB and DAEO. The first version of the manuscript was written by DAEO, and all authors commented on it. All authors read and agreed on the final version of the manuscript.

### Data availability

The moss vouchers are deposited at the QFA herbarium corresponding to catalogue numbers QFA-637608 to QFA-637686. The raw sequences of this study are deposited in the NCBI Sequence Read Archive (SRA) database under the BioProject PRJNA893897. The moss and soil samples are associated with BioSamples SAMN31436785–SAMN31436859 and SAMN31439125– SAMN31439151, respectively. The moss 16S sequences correspond to SRA accessions SRR22028313–SRR22028356, moss *nif*H sequences to SRR22032306– SRR22032359, and soil 16S sequences to SRR22031903– SRR22031923. For detailed information, see Supplementary Information Table S1. The dataset and scripts for bioinformatic analyses are available from the corresponding author upon request.

## REFERENCES

Ainon H., Tan CJ, Vikineswary S (2006) Biological Characterization of Rhodomicrobium vannielii Isolated from a Hot Spring at Gadek, Malacca, Malaysia. Malays J Microbiol. https://doi.org/10.21161/mjm.210603

Alcaraz LD, Peimbert M, Barajas HR, et al (2018) Marchantia liverworts as a proxy to plants’ basal microbiomes. Sci Rep 8:1–12. https://doi.org/10.1038/s41598-018-31168-0

Alonso-García M, Villarreal Aguilar JC (2022) Bacterial community of reindeer lichens differs between northern and southern lichen woodlands. Can J For Res 12:1–12. https://doi.org/10.1139/cjfr-2021-0272

Alvarenga DO, Rousk K (2021) Indirect effects of climate change inhibit N2 fixation associated with the feathermoss Hylocomium splendens in subarctic tundra. Sci Total Environ 795:148676. https://doi.org/10.1016/j.scitotenv.2021.148676

Alvarenga DO, Rousk K (2022) Unraveling host-microbe interactions and ecosystem functions in moss-bacteria symbioses. J. Exp. Bot. 73:4473–4486

Angel R, Nepel M, Panhölzl C, et al (2018) Evaluation of Primers Targeting the Diazotroph Functional Gene and Development of NifMAP – A Bioinformatics Pipeline for Analyzing nifH Amplicon Data. Front Microbiol 9:1–15. https://doi.org/10.3389/fmicb.2 018.00703

Bay G, Nahar N, Oubre M, et al (2013a) Boreal feather mosses secrete chemical signals to gain nitrogen. New Phytol 200:54–60. https://doi.org/10.1111/nph.12403

Bay G, Nahar N, Oubre M, et al (2013b) Boreal feather mosses secrete chemical signals to gain nitrogen. New Phytol 200:54–60. https://doi.org/10.1111/nph.12403

Bell-Doyon P, Bellavance V, Bélanger L, et al (2022) Bacterial, Fungal, and Mycorrhizal Communities in the Soil Differ between Clearcuts and Insect Outbreaks in the Boreal Forest 50 Years after Disturbance. SSRN Electron J 4:1–55. https://doi.org/10.2139/ssrn.4140847

Bellenger JP, Darnajoux R, Zhang X, Kraepiel AML (2020) Biological nitrogen fixation by alternative nitrogenases in terrestrial ecosystems: a review. Biogeochemistry 149:53–73. https://doi.org/10.1007/s10533-020-00666-7

Bouchard R, Peñaloza-Bojacá G, Toupin S, et al (2020) Contrasting bacteriome of the hornwort Leiosporoceros dussii in two nearby sites with emphasis on the hornwort-cyanobacterial symbiosis. Symbiosis 81:39–52. https://doi.org/10.1007/s13199-020-00680-1

Bragina A, Berg C, Berg G (2015) The core microbiome bonds the Alpine bog vegetation to a transkingdom metacommunity. Mol Ecol 24:4795–4807. https://doi.org/10.1111/mec.13342

Brooks, Mollie E, Kristensen K, Benthem, Koen, J. V, et al (2017) glmmTMB Balances Speed and Flexibility Among Packages for Zero-inflated Generalized Linear Mixed Modeling. R J 9:378. https://doi.org/10.32614/RJ-2017-066

Calabria LM, Petersen KS, Bidwell A, Hamman ST (2020) Moss-cyanobacteria associations as a novel source of biological N2-fixation in temperate grasslands. Plant Soil 456:307–321. https://doi.org/10.1007/s11104-020-04695-x

Callahan B (2018) Silva taxonomic training data formatted for DADA2 (Silva version 132)

Callahan BJ, McMurdie PJ, Holmes SP (2017) Exact sequence variants should replace operational taxonomic units in marker-gene data analysis. ISME J 11:2639–2643. https://doi.org/10.1038/ismej.2017.119

Callahan BJ, McMurdie PJ, Rosen MJ, et al (2016) DADA2: High-resolution sample inference from Illumina amplicon data. Nat Methods 13:581–583. https://doi.org/10.1038/nmeth.3869

Camacho C, Coulouris G, Avagyan V, et al (2009) BLAST+: Architecture and applications. BMC Bioinformatics 10:1–9. https://doi.org/10.1186/1471-2105-10-421/FIGURES/4

Carella P, Schornack S (2018) Manipulation of Bryophyte Hosts by Pathogenic and Symbiotic Microbes Special Issue - Review. 0:1–10. https://doi.org/10.1093/pcp/pcx182

Carrell AA, Kolton M, Glass JB, et al (2019) Experimental warming alters the community composition, diversity, and N2 fixation activity of peat moss (Sphagnum fallax) microbiomes. Glob Chang Biol 25:2993–3004. https://doi.org/10.1111/gcb.14715

Chapin III FS, Jefferies RL, Reynolds JF, et al (1992) Arctic Ecosystems in a Changing Climate, 1st editio. Academic Press. Elsevier

Chen J, Bittinger K, Charlson ES, et al (2012) Associating microbiome composition with environmental covariates using generalized UniFrac distances. Bioinformatics 28:2106– 2113. https://doi.org/10.1093/bioinformatics/bts342

Choudhary S, Blaud A, Osborn AM, et al (2016) Nitrogen accumulation and partitioning in a High Arctic tundra ecosystem from extreme atmospheric N deposition events. Sci Total Environ 554–555:303–310. https://doi.org/10.1016/j.scitotenv.2016.02.155

Cutler N (2011) Nutrient limitation during long-term ecosystem development inferred from a mat-forming moss. Bryologist 114:204–214. https://doi.org/10.1639/0007-2745.114.1.204

Daniëls FJ a, Gillespie LJ, Poulin M, et al (2013) Plants

Darnajoux R, Magain N, Renaudin M, et al (2019) Molybdenum threshold for ecosystem scale alternative vanadium nitrogenase activity in boreal forests. Proc Natl Acad Sci 116:201913314. https://doi.org/10.1073/pnas.1913314116

Davis NM, Proctor DiM, Holmes SP, et al (2018) Simple statistical identification and removal of contaminant sequences in marker-gene and metagenomics data. Microbiome 6:1–14. https://doi.org/10.1186/S40168-018-0605-2/FIGURES/6

de Jesús Suárez-Moo P, Vovides AP, Patrick Griffith M, et al (2019) Unlocking a high bacterial diversity in the coralloid root microbiome from the cycad genus Dioon. PLoS One 14:1–20. https://doi.org/10.1371/journal.pone.0211271

DeLuca TH, Zackrisson O, Nilsson MC, Sellstedt A (2002) Quantifying nitrogen-fixation in feather moss carpets of boreal forests. Nature 419:917–920. https://doi.org/10.1038/nature01051

Eddy SR (2011) Accelerated Profile HMM Searches. PLOS Comput Biol 7:e1002195. https://doi.org/10.1371/JOURNAL.PCBI.1002195

Edgar RC, Bateman A (2010) Search and clustering orders of magnitude faster than BLAST. Bioinformatics 26:2460–2461. https://doi.org/10.1093/BIOINFORMATICS/BTQ461

Escolástico-Ortiz DA, Hedenäs L, Quandt D, et al (2022) Cryptic speciation shapes the biogeographic history of a northern distributed moss. Bot J Linn Soc 1–21. https://doi.org/10.1093/botlinnean/boac027

Faith DP (1992) Conservation evaluation and phylogenetic diversity. Biol Conserv 61:1–10. https://doi.org/10.1016/0006-3207(92)91201-3

Frank IE, Turk-Kubo KA, Zehr JP (2016) Rapid annotation of nifH gene sequences using classification and regression trees facilitates environmental functional gene analysis. Environ Microbiol Rep 8:905–916. https://doi.org/10.1111/1758-2229.12455

Gentili F, Nilsson MC, Zackrisson O, et al (2005) Physiological and molecular diversity of feather moss associative N 2-fixing cyanobacteria. J Exp Bot 56:3121–3127. https://doi.org/10.1093/jxb/eri309

Gordon C, Wynn JM, Woodin SJ (2001) Impacts of increased nitrogen supply on high Arctic heath: The importance of bryophytes and phosphorus availability. New Phytol 149:461–471. https://doi.org/10.1046/j.1469-8137.2001.00053.x

Hardy RWF, Holsten RD, Jackson EK, Burns RC (1968) The Acetylene - Ethylene Assay for N2 FixationLJ: Laboratory and Field Evaluation. Plant Physiol 43:1185–1207

Hare FK, Ritchie JC (1972) The Boreal Bioclimates. Geogr Rev 62:333. https://doi.org/10.2307/213287

Harwood CS (2020) Iron-Only and Vanadium Nitrogenases: Fail-Safe Enzymes or Something More? Annu Rev Microbiol 74:247–266. https://doi.org/10.1146/annurev-micro-022620-014338

Hedenäs L (2020) Cryptic speciation revealed in Scandinavian Racomitrium lanuginosum (Hedw.) Brid. (Grimmiaceae). J Bryol 42:117–127. https://doi.org/10.1080/03736687.2020.1722923

Hodgetts N (2019) Racomitrium lanuginosum. In: IUCN Red List Threat. Species 2019. https://www.iucnredlist.org/species/85844273/87795587

Holland-Moritz H, Stuart J, Lewis LR, et al (2018) Novel bacterial lineages associated with boreal moss species. Environ Microbiol 20:2625–2638. https://doi.org/10.1111/1462-2920.14288

Holland-Moritz H, Stuart JEM, Lewis LR, et al (2021) The bacterial communities of Alaskan mosses and their contributions to N2-fixation. Microbiome 9:53. https://doi.org/10.1186/s40168-021-01001-4

Jean M, Fenton NJ, Bergeron Y, Nilsson MC (2021) Sphagnum and feather moss-associated N2 fixation along a 724-year chronosequence in eastern boreal Canada. Plant Ecol 222:1007– 1022. https://doi.org/10.1007/s11258-021-01157-x

Jean M, Holland-Moritz H, Melvin AM, et al (2020) Experimental assessment of tree canopy and leaf litter controls on the microbiome and nitrogen fixation rates of two boreal mosses. New Phytol 227:1335–1349. https://doi.org/10.1111/nph.16611

Kembel SW, Cowan PD, Helmus MR, et al (2010) Picante: R tools for integrating phylogenies and ecology. Bioinformatics 26:1463–1464. https://doi.org/10.1093/bioinformatics/btq166

Klarenberg IJ, Keuschnig C, Russi Colmenares AJ, et al (2021) LongLJterm warming effects on the microbiome and nifH gene abundance of a common moss species in subLJArctic tundra. New Phytol. https://doi.org/10.1111/nph.17837

Klarenberg IJ, Keuschnig C, Salazar A, et al (2023) Moss and underlying soil bacterial community structures are linked to moss functional traits. Ecosphere 14:1–16. https://doi.org/10.1002/ecs2.4447

Kostka JE, Weston DJ, Glass JB, et al (2016) The Sphagnum microbiome: New insights from an ancient plant lineage. New Phytol 211:57–64. https://doi.org/10.1111/nph.13993

Kulichevskaya IS, Ivanova AO, Baulina OI, et al (2008) Singulisphaera acidiphila gen. nov., sp. nov., a non-filamentous, Isosphaera-like planctomycete from acidic northen wetlands. Int J Syst Evol Microbiol 58:1186–1193. https://doi.org/10.1099/ijs.0.65593-0

Kulichevskaya IS, Suzina NE, Liesack W, Dedysh SN (2010) Bryobacter aggregatus gen. nov., sp. nov., a peat-inhabiting, aerobic chemo-organotroph from subdivision 3 of the acidobacteria. Int J Syst Evol Microbiol 60:301–306. https://doi.org/10.1099/ijs.0.013250-0

Lahti L, Shetty S (2017) Tools for microbiome analysis in R

Lajoie G, Kembel SW (2021) Host neighborhood shapes bacterial community assembly and specialization on tree species across a latitudinal gradient. Ecol Monogr 91:. https://doi.org/10.1002/ecm.1443

Larraín J, Quandt D, Stech M, Muñoz J (2013) Lumping or splitting? The case of Racomitrium (Bryophytina: Grimmiaceae). Taxon 62:1117–1132. https://doi.org/10.12705/626.45

Larsson J (2022) eulerr: Area-Proportional Euler and Venn Diagrams with Ellipses

Lee KB, De Backer P, Aono T, et al (2008) The genome of the versatile nitrogen fixer Azorhizobium caulinodans ORS571. BMC Genomics 9:1–14. https://doi.org/10.1186/1471-2164-9-271

Leppänen SM, Salemaa M, Smolander A, et al (2013) Nitrogen fixation and methanotrophy in forest mosses along a N deposition gradient. Environ Exp Bot 90:62–69. https://doi.org/10.1016/j.envexpbot.2012.12.006

Liu X, Rousk K (2022) The moss traits that rule cyanobacterial colonization. Ann Bot 129:147–160. https://doi.org/10.1093/aob/mcab127

Madigan M, Cox SS, Stegeman RA (1984) Nitrogen fixation and nitrogenase activities in members of the family Rhodospirillaceae. J Bacteriol 157:73–78. https://doi.org/10.1128/jb.157.1.73-78.1984

McMurdie PJ, Holmes S (2013) phyloseq: An R Package for Reproducible Interactive Analysis and Graphics of Microbiome Census Data. PLoS One 8:e61217. https://doi.org/10.1371/journal.pone.0061217

Morrow MA, Pold G, DeAngelis KM (2020) Draft Genome Sequence of a Terrestrial Planctomycete, Singulisphaera sp. Strain GP187, Isolated from Forest Soil. Microbiol Resour Announc 9:18–20. https://doi.org/10.1128/MRA.00956-20

Moyes AB, Kueppers LM, Pett-Ridge J, et al (2016) Evidence for foliar endophytic nitrogen fixation in a widely distributed subalpine conifer. New Phytol 210:657–668. https://doi.org/10.1111/nph.13850

Murray MG, Thompson WF (1980) Research Rapid isolation of high molecular weight plant DNA. Nucleic Acids 8:4321–4325

Oksanen J, Blanchet FG, Friendly M, et al (2018) vegan: Community Ecology Package Payette S (1993) The range limit of boreal tree species in Québec-Labrador: an ecological and palaeoecological interpretation. Rev Palaeobot Palynol 79:7–30. https://doi.org/10.1016/0034-6667(93)90036-T

Payette S, Fortin M-J, Gamache I (2001) The Subarctic Forest–Tundra: The Structure of a Biome in a Changing Climate. Bioscience 51:709–718. https://doi.org/https://doi.org/10.1641/0006-3568(2001)051[0709:TSFTTS]2.0.CO;2

Prairie Climate Centre, University of Winnipeg (2022) Climate Atlas of Canada. In: Mean Temp. (Annual). Munic. Kuujjuarapik. https://climateatlas.ca/map/canada/annual_meantemp_2030_85#z=8&lat=54.88&lng=-77.47&city=25. Accessed 30 Nov 2022

Pruitt KD, Tatusova T, Maglott DR (2005) NCBI Reference Sequence (RefSeq): a curated non-redundant sequence database of genomes, transcripts and proteins. Nucleic Acids Res 33:D501–D504. https://doi.org/10.1093/NAR/GKI025

R Core Team (2017) R Development Core Team. R A Lang. Environ. Stat. Comput. 55:275–286

Renaudin M, Blasi C, Bradley RL, Bellenger J (2022a) New insights into the drivers of mossLJassociated nitrogen fixation and cyanobacterial biomass in the eastern Canadian boreal forest. J Ecol 102:1–3. https://doi.org/10.1111/1365-2745.13881

Renaudin M, Laforest-Lapointe I, Bellenger J (2022b) Unraveling global and diazotrophic bacteriomes of boreal forest floor feather mosses and their environmental drivers at the ecosystem and at the plant scale in North America. Sci Total Environ 837:155761. https://doi.org/10.1016/j.scitotenv.2022.155761

RodríguezLJRodríguez JC, Bergeron Y, Kembel SW, Fenton NJ (2022) Dominance of coniferous and broadleaved trees drives bacterial associations with boreal feather mosses. Environ Microbiol 24:3517–3528. https://doi.org/10.1111/1462-2920.16013

Rousk K (2022) Biotic and abiotic controls of nitrogen fixation in cyanobacteria–moss associations. New Phytol 1330–1335. https://doi.org/10.1111/nph.18264

Rousk K, Degboe J, Michelsen A, et al (2017a) Molybdenum and phosphorus limitation of moss-associated nitrogen fixation in boreal ecosystems. New Phytol 214:97–107. https://doi.org/10.1111/nph.14331

Rousk K, Jones DL, DeLuca TH (2013) Moss-cyanobacteria associations as biogenic sources of nitrogen in boreal forest ecosystems. Front Microbiol 4:1–10. https://doi.org/10.3389/fmicb.2013.00150

Rousk K, Rousk J (2020) The responses of moss-associated nitrogen fixation and belowground microbial community to chronic Mo and P supplements in subarctic dry heaths. Plant Soil 451:261–276. https://doi.org/10.1007/s11104-020-04492-6

Rousk K, Sorensen PL, Michelsen A (2017b) Nitrogen fixation in the High Arctic: a source of ‘new’ nitrogen? Biogeochemistry 136:213–222. https://doi.org/10.1007/s10533-017-0393-y

Rzepczynska AM, Michelsen A, Olsen MAN, Lett S (2022) Bryophyte species differ widely in their growth and N 2 -fixation responses to temperature. Arct Sci. https://doi.org/10.1139/as-2021-0053

Schliep KP (2011) phangorn: phylogenetic analysis in R. Bioinformatics 27:592–593. https://doi.org/10.1093/bioinformatics/btq706

Segata N, Izard J, Waldron L, et al (2011) Metagenomic biomarker discovery and explanation. Genome Biol 12:R60. https://doi.org/10.1186/GB-2011-12-6-R60

Shade A, Handelsman J (2012) Beyond the Venn diagram: the hunt for a core microbiome. Environ Microbiol 14:4–12. https://doi.org/10.1111/j.1462-2920.2011.02585.x

Stamatakis A (2014) RAxML version 8: A tool for phylogenetic analysis and post-analysis of large phylogenies. Bioinformatics 30:1312–1313. https://doi.org/10.1093/bioinformatics/btu033

Stech M, Veldman S, Larraín J, et al (2013) Molecular Species Delimitation in the Racomitrium canescens Complex (Grimmiaceae) and Implications for DNA Barcoding of Species Complexes in Mosses. PLoS One 8:. https://doi.org/10.1371/journal.pone.0053134

Stuart JEM, Holland-Moritz H, Lewis LR, et al (2021) Host Identity as a Driver of Moss-Associated N2 Fixation Rates in Alaska. Ecosystems 24:530–547. https://doi.org/10.1007/s10021-020-00534-3

Ullrich SR, Poehlein A, Voget S, et al (2015) Permanent draft genome sequence of Acidiphilium sp. JA12–A1. Stand Genomic Sci 10:. https://doi.org/10.1186/s40793-015-0040-y

Villarreal A. JC, Renaudin M, Beaulieu-Laliberté A, Bellenger JP (2021) Stigonema associated with boreal Stereocaulon possesses the alternative vanadium nitrogenase. Lichenol 53:215–220. https://doi.org/10.1017/S0024282921000062

Vitt DH, Marsh C (1988) Population variation and phytogeography of Racomitrium lanuginosum and R. pruinosum. Beih zur Nov Hedwigia 90:235–260

Wang Q, Quensen JF, Fish JA, et al (2013) Ecological patterns of nifH genes in four terrestrial climatic zones explored with targeted metagenomics using framebot, a new informatics tool. MBio 4:. https://doi.org/10.1128/MBIO.00592-13/SUPPL_FILE/MBO005131620S1.PDF

Ward LM, Cardona T, Holland-Moritz H (2019) Evolutionary Implications of Anoxygenic Phototrophy in the Bacterial Phylum Candidatus Eremiobacterota (WPS-2). Front Microbiol 10:1–12. https://doi.org/10.3389/fmicb.2019.01658

Warshan D, Bay G, Nahar N, et al (2016) Seasonal variation in nifH abundance and expression of cyanobacterial communities associated with boreal feather mosses. ISME J 10:2198– 2208. https://doi.org/10.1038/ismej.2016.17

Warshan D, Espinoza JL, Stuart RK, et al (2017) Feathermoss and epiphytic Nostoc cooperate differently: Expanding the spectrum of plant-cyanobacteria symbiosis. ISME J 11:2821– 2833. https://doi.org/10.1038/ismej.2017.134

Wright, Erik S (2016) Using DECIPHER v2.0 to Analyze Big Biological Sequence Data in R. R J 8:352. https://doi.org/10.32614/RJ-2016-025

Zhang X, McRose DL, Darnajoux R, et al (2016) Alternative nitrogenase activity in the environment and nitrogen cycle implications. Biogeochemistry 127:189–198. https://doi.org/10.1007/s10533-016-0188-6

