## Supplementary Information - Figures for "Differentially abundant bacteria drive the N_2_-fixation of a widespread moss in the forest-tundra transition zone"

Symbiosis

Dennis Alejandro Escolástico-Ortiz[1,2,3*], Charlotte Blasi[4], Jean-Philippe Bellenger[4], Nicolas Derome[2] and Juan Carlos Villarreal[1,2,3]

[1]Département de Biologie, Université Laval, Québec, G1V 0A6, Canada
[2]Institut de Biologie Intégrative et des Systèmes (IBIS), Université Laval, Québec, G1V 0A6, Canada.
[3]Centre d’études nordiques (CEN), Université Laval, Québec, QC, Canada
[4]Centre Sève, Département de Chimie, Université de Sherbrooke, Sherbrooke, J1K 2R1, QC, Canada
* ****

### Supplementary Information

### Figures

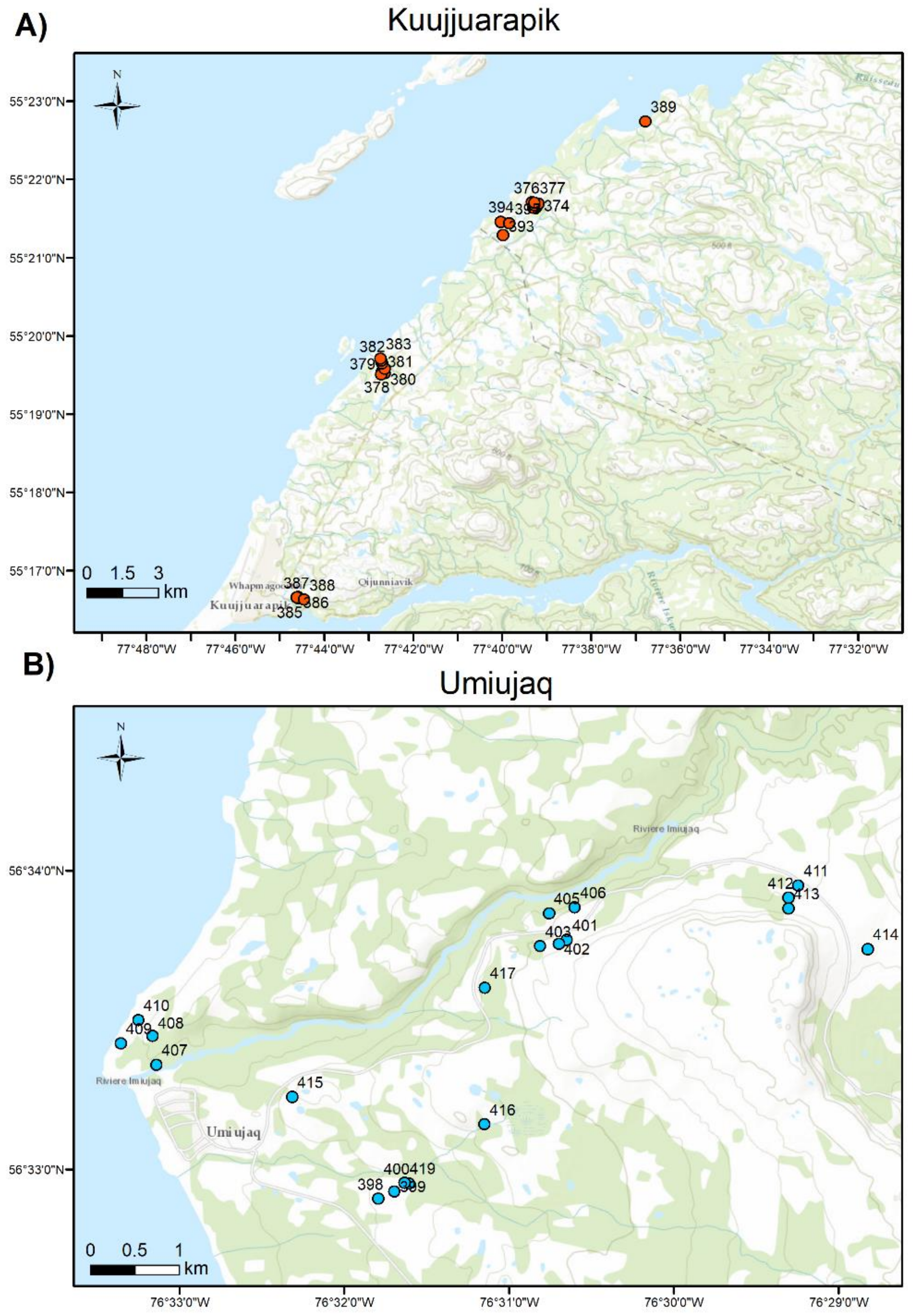


**Fig. S1** Populations and samples of *Racomitrium lanuginosum* collected for this study. A) Kuujjuarapik population indicating the location of each sampling plot in red (n =15). The vegetation here is forest tundra. B) Umiujaq population representing each plot in blue (n = 12). This area is characterized by shrub tundra. Note that scales are different for each map, and the final dataset did not include all plots for analyses

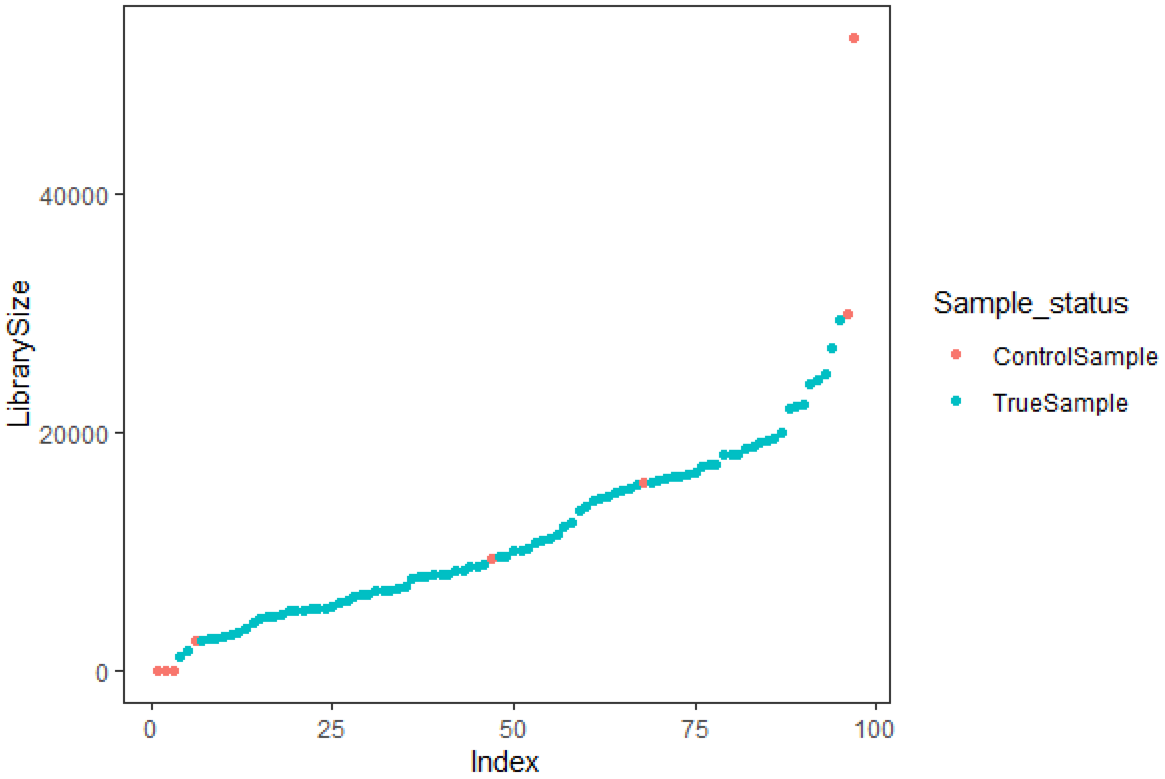


**Fig. S2** Sample library sizes based on 16S rRNA gene amplicons of *Racomitrium lanuginosum* and associated soil. Library size is presented as the number of reads per sample. Index on the y-axis refers to the position of samples according to the number of reads. The sample status is indicated for those used in analyses in blue (True sample) and sequenced negative control PCRs in red (Control sample). Control samples are expected to have a small library size; however, one control sample has a higher-than-expected library size

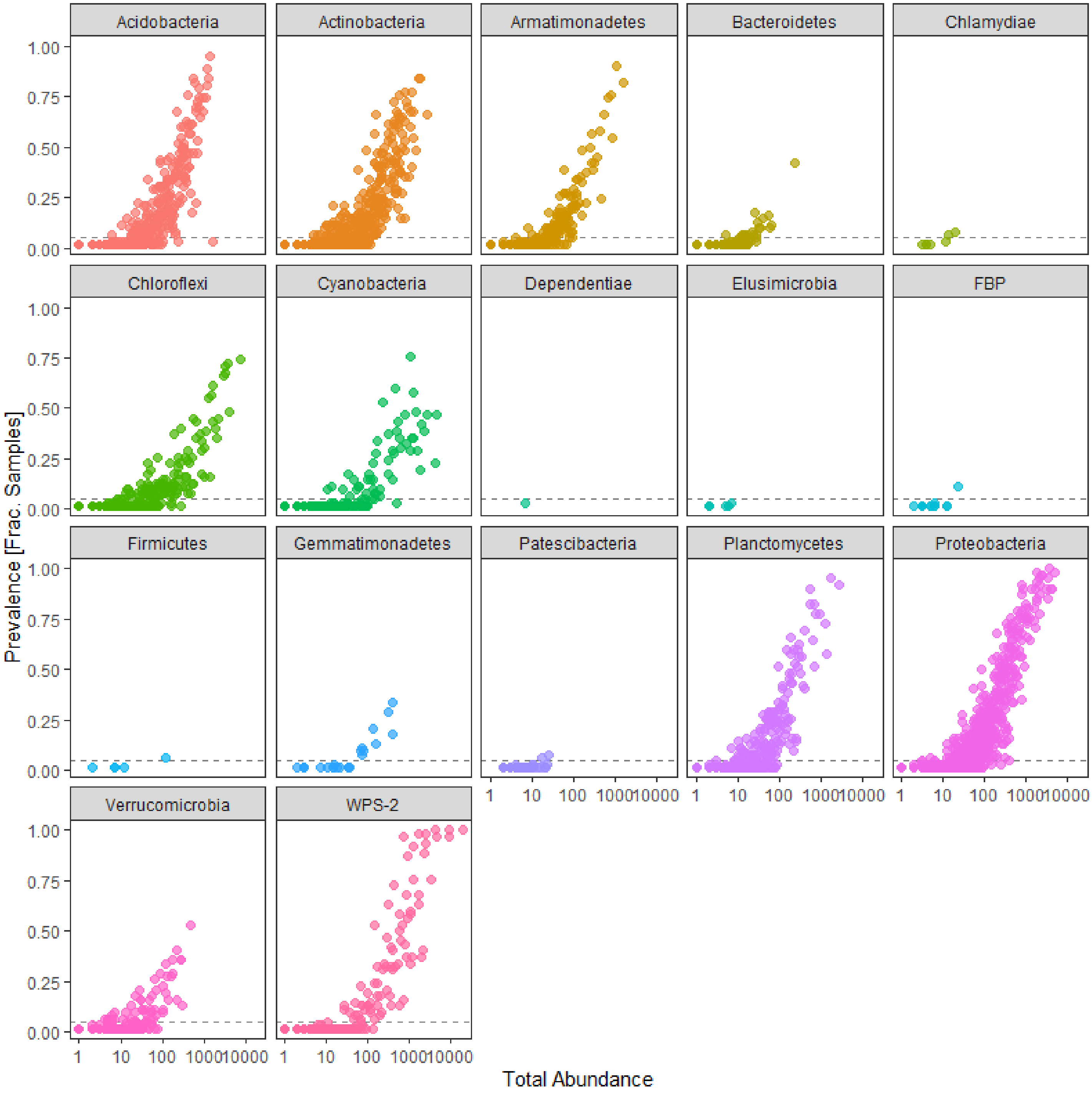


**Fig. S3** Prevalence and total abundance of ASVs per phylum based on *Racomitrium lanuginosum* 16S rRNA gene. The proportion of samples where an ASV occurs is presented on the y-axis, and total abundance is on the x-axis. The dotted line indicates the 0.05 prevalence threshold used to filter out ASVs occurring in less than three samples. Acidobacteria, Actinobacteria, Planctomycetes, Proteobacteria and WPS-2 (Eremiobacterota) have a high prevalence and abundance

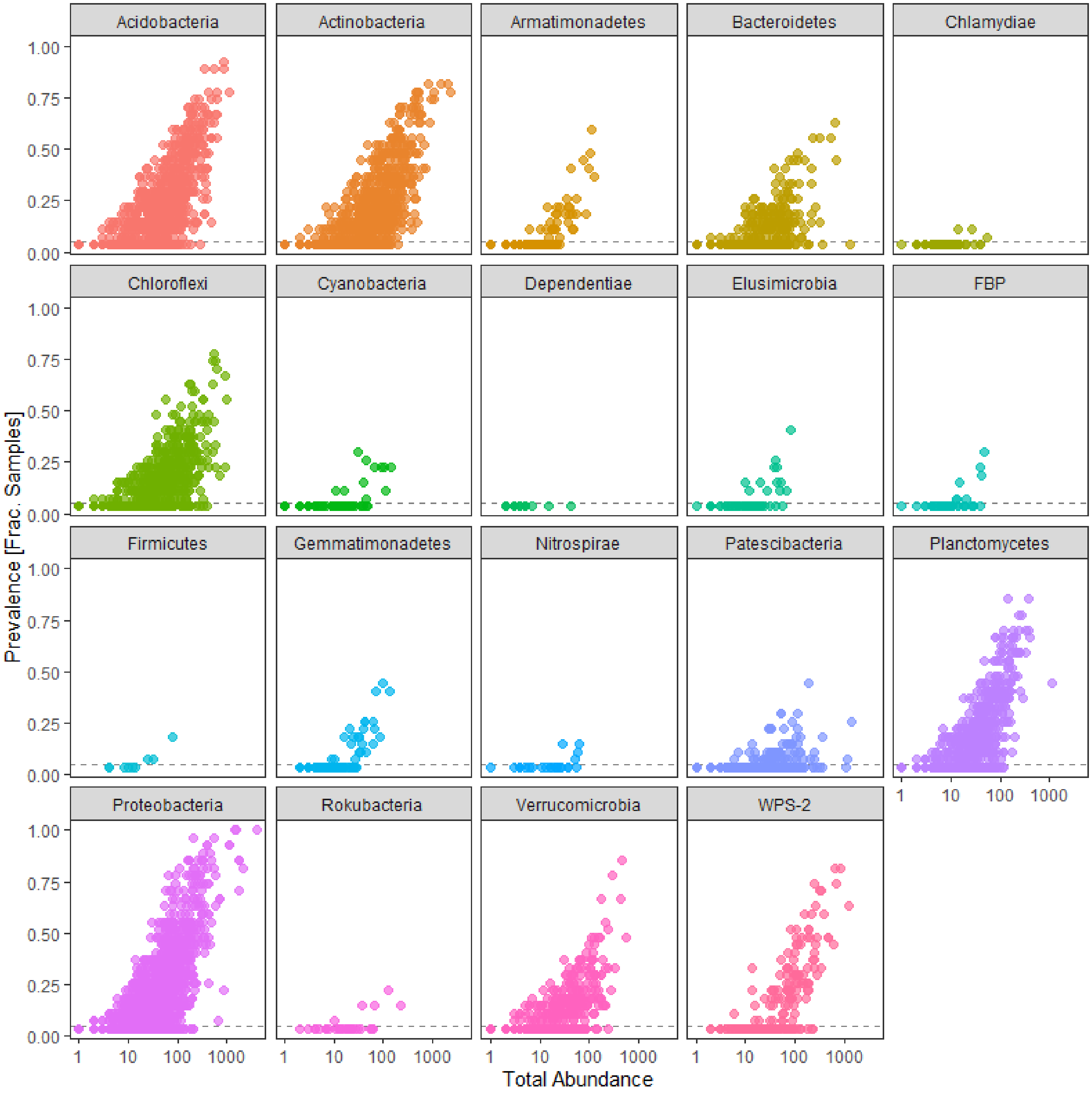


**Fig. S4** Prevalence and total abundance of ASVs per phylum based on 16S rRNA gene of soil samples associated with *Racomitrium lanuginosum*. The proportion of samples where an ASV occurs is presented on the y-axis and total abundance is on the x-axis. The dotted line indicates the 0.05 prevalence threshold used to filter out ASVs occurring in less than one sample. Acidobacteria, Actinobacteria, Planctomycetes and Proteobacteria have a high prevalence and abundance

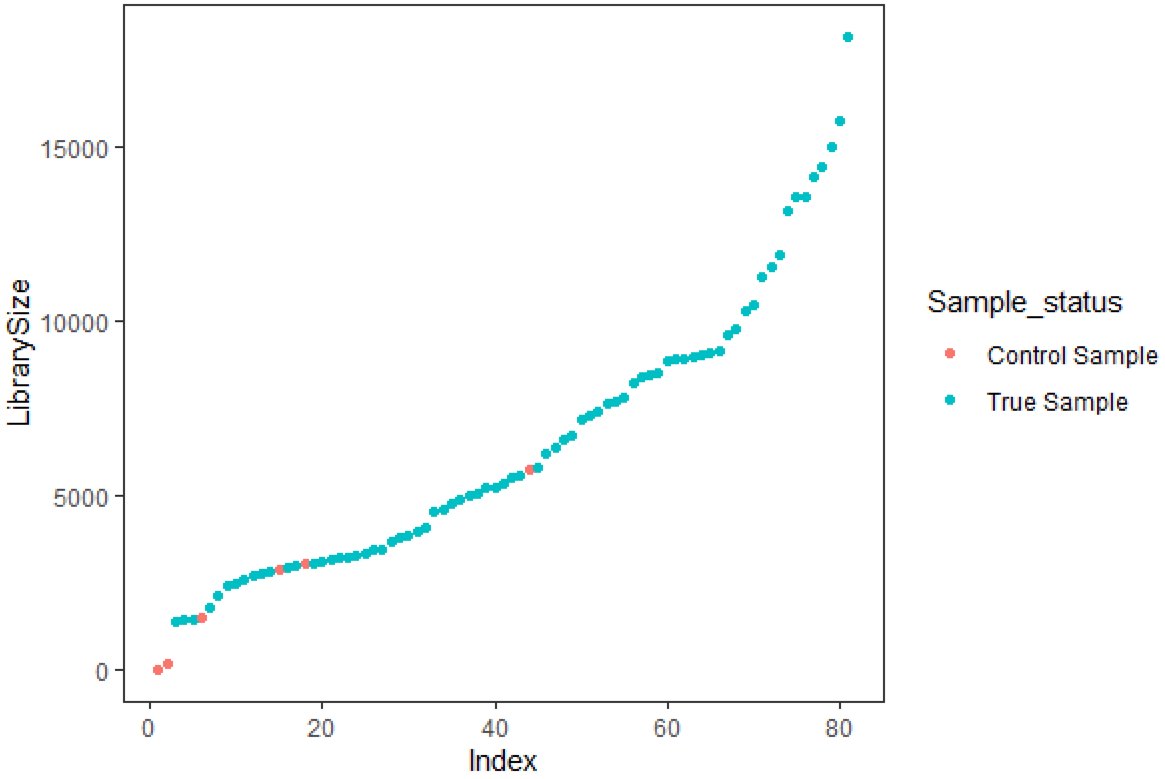


**Fig. S5** Sample library sizes based on *nif*H gene amplicons. Library size is presented as the number of reads per sample. Index on the y-axis refers to the position of samples according to the number of reads. The sample status is indicated for those used in analyses in blue (True sample) and sequenced negative control PCRs in red (Control sample). Control samples have a smaller library size compared with true samples

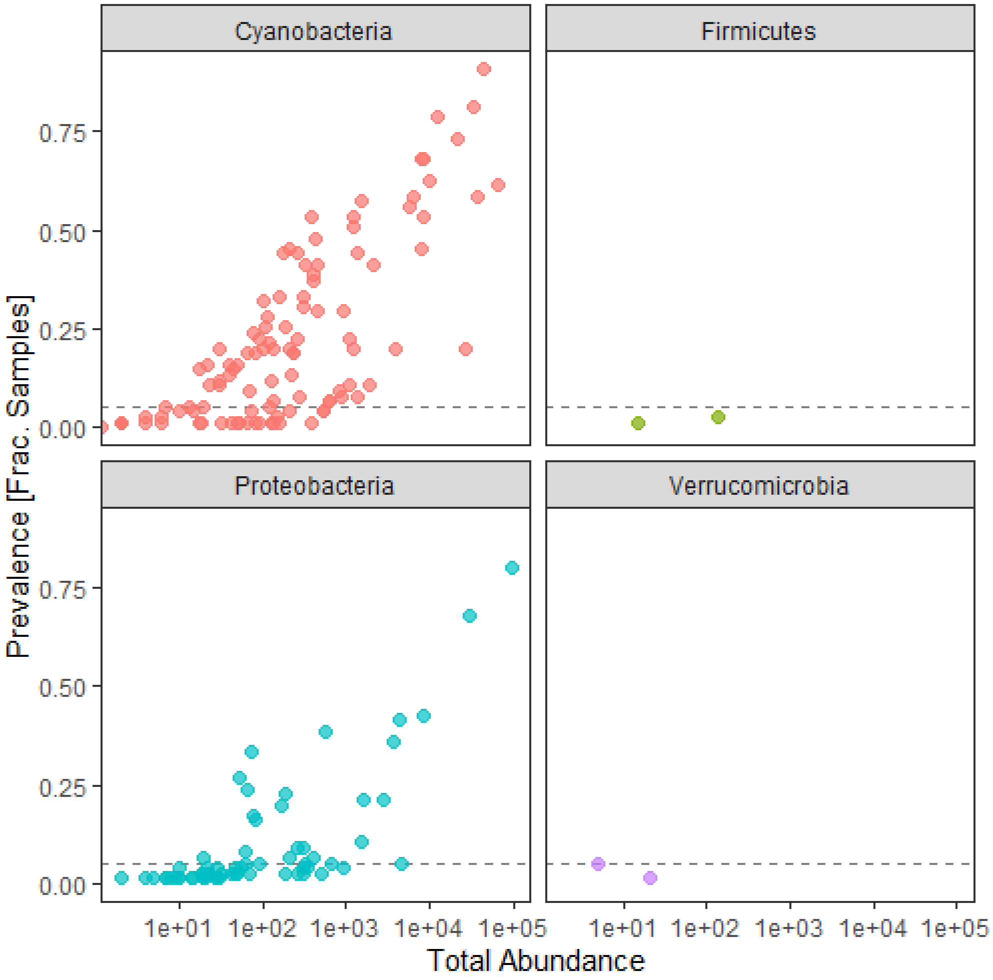


**Fig. S6** Prevalence and total abundance of OTUs per phylum based on *nif*H gene. The proportion of samples where an OTU occurs is presented on the y-axis, and total abundance is on the x-axis. The dotted line indicates the 0.05 prevalence threshold used to filter out OTUs occurring only in three samples. Cyanobacteria and Proteobacteria dominate the diazotrophic community

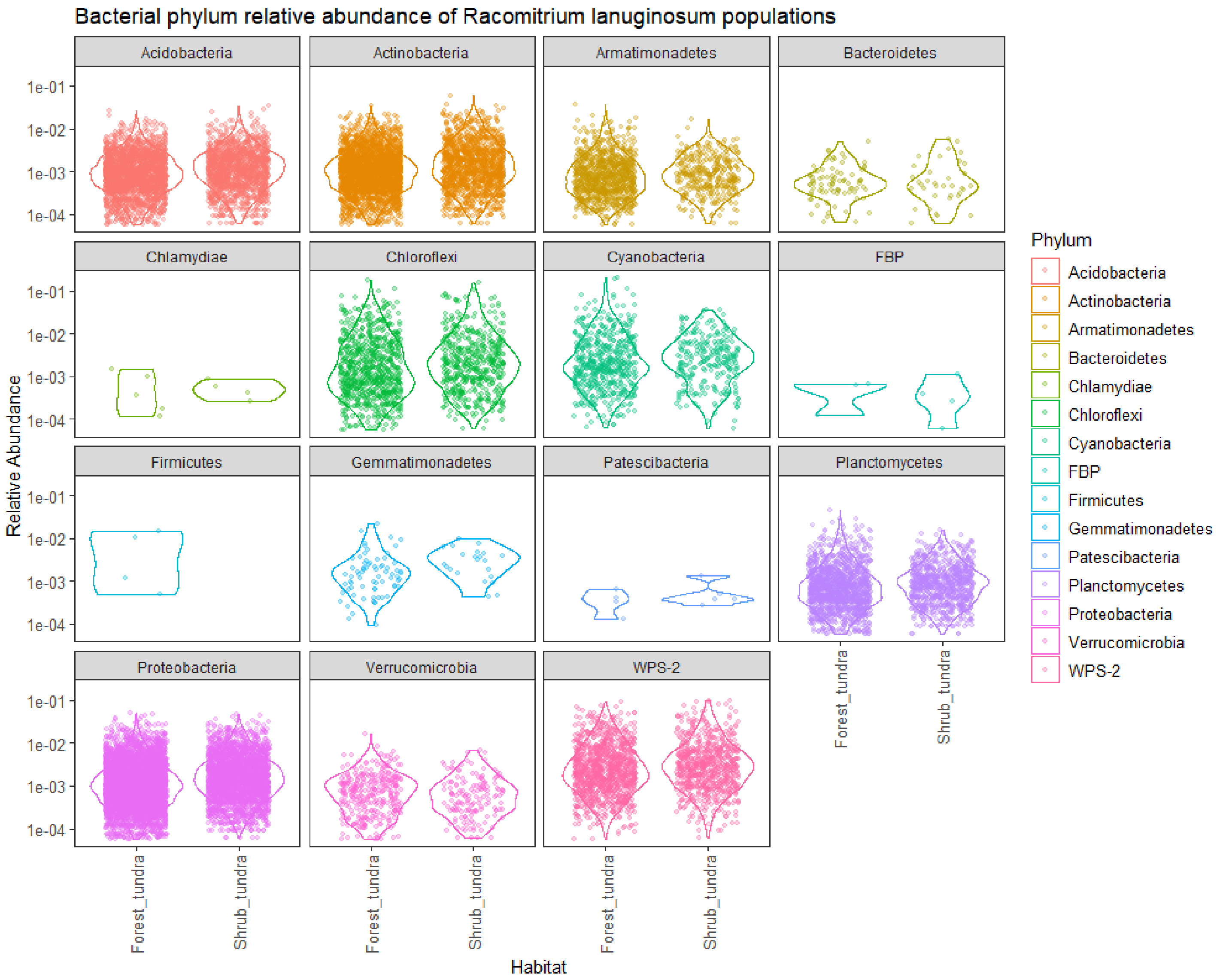


**Fig. S7** Relative abundance per phylum of bacterial ASVs of two *Racomitrium lanuginosum* populations based on 16S rRNA data. The most abundant phyla are Acidobacteria, Actinobacteria, Chloroflexi, Proteobacteria and WPS-2 (Eremiobacterota). The ASV relative abundance between the forest and shrub tundra does not seem to differ

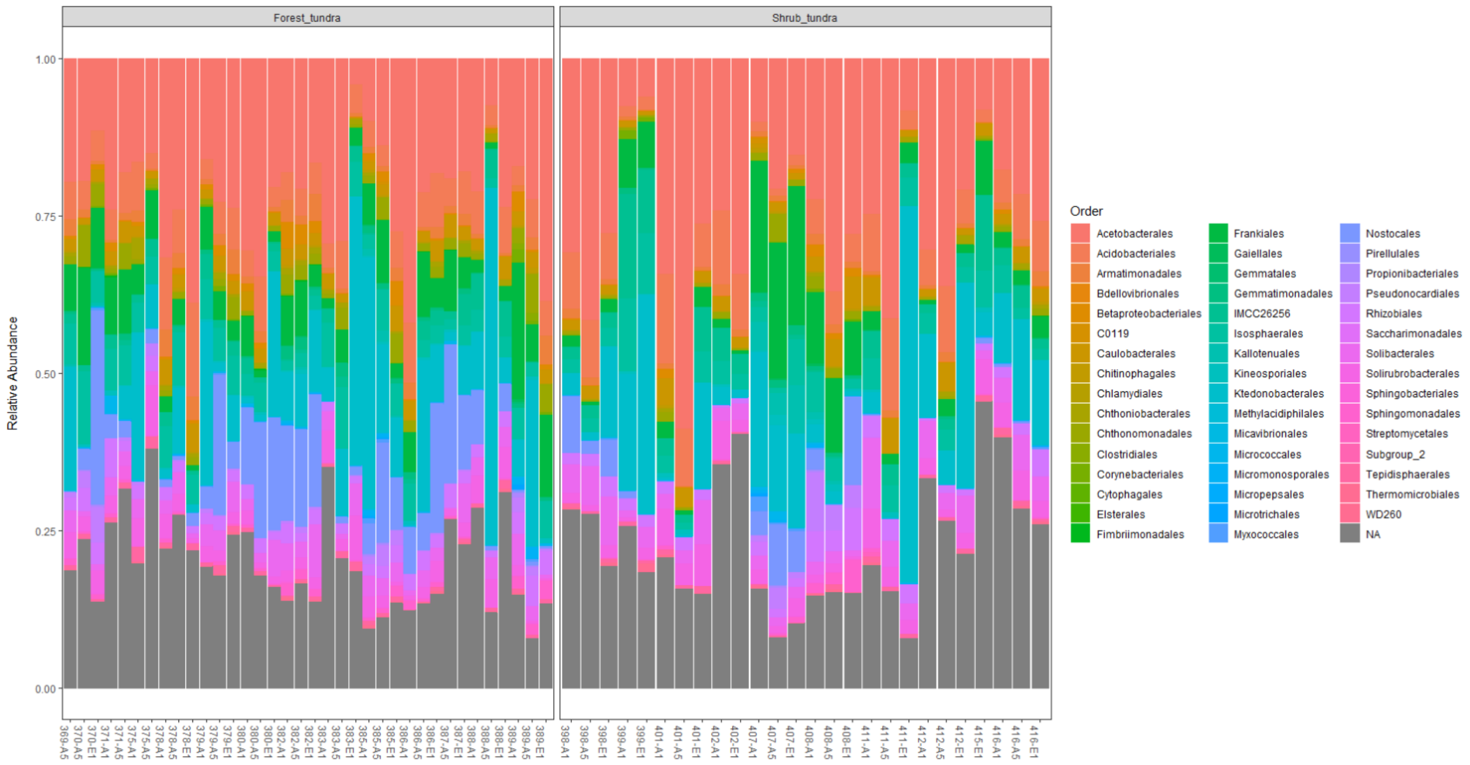


**Fig. S8** Relative abundance per order of bacterial ASVs of two *Racomitrium lanuginosum* populations based on 16S rRNA data. The most abundant orders are Acetobacterales, Acidobacterales, Isosphaerales, Ktedonobacterales and Frankiales. ASV abundances seem similar in the forest tundra and shrub tundra

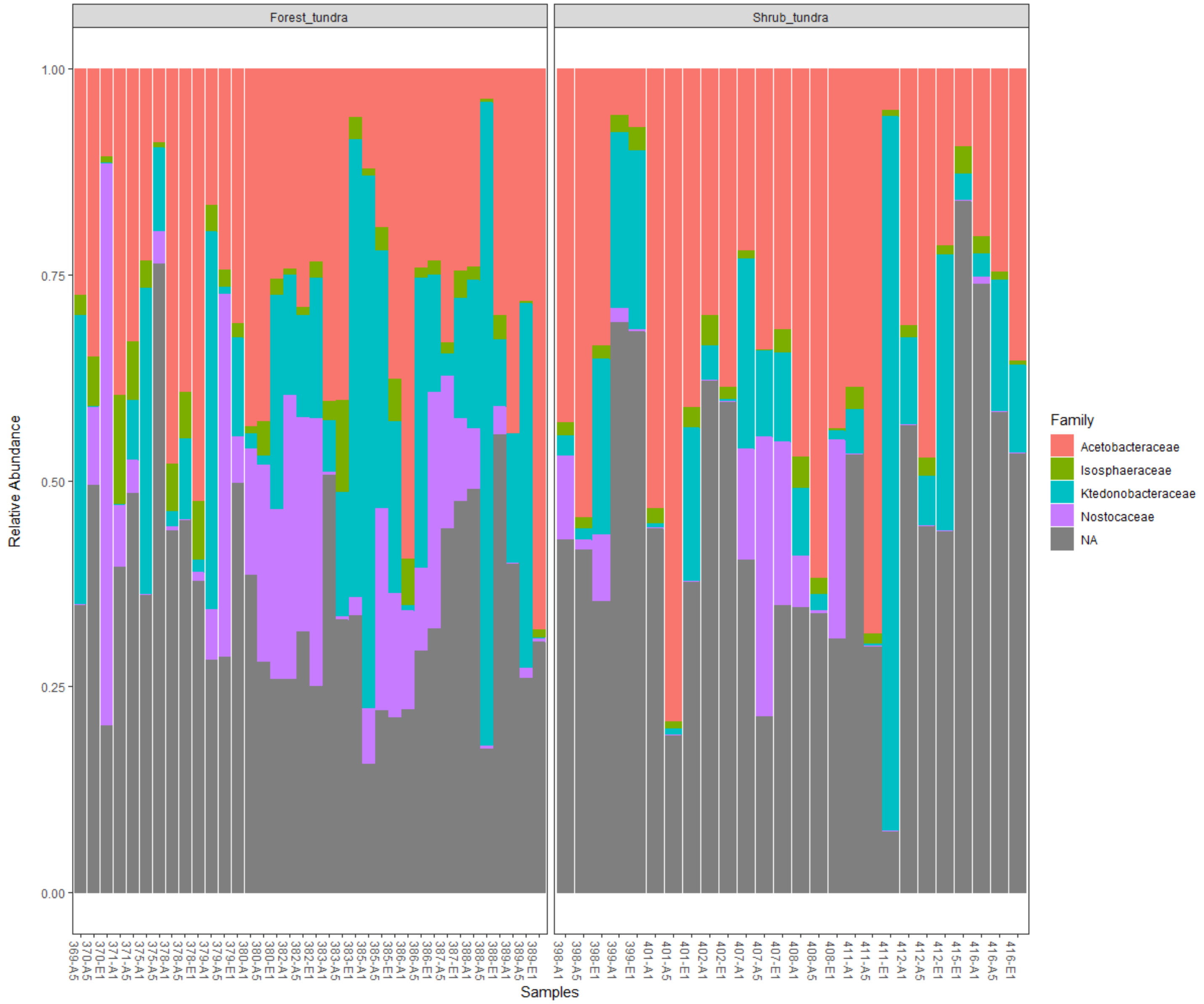


**Fig. S9** Relative abundance per family of the 50 most abundant bacterial ASVs of two *Racomitrium lanuginosum* populations based on 16S rRNA data. Dominant ASVs were mainly represented by Acetobacteraceae, Eremiobacterota unclassified group and Ktedonobacteraceae. Nostocaceae seem to be better represented in the forest-tundra

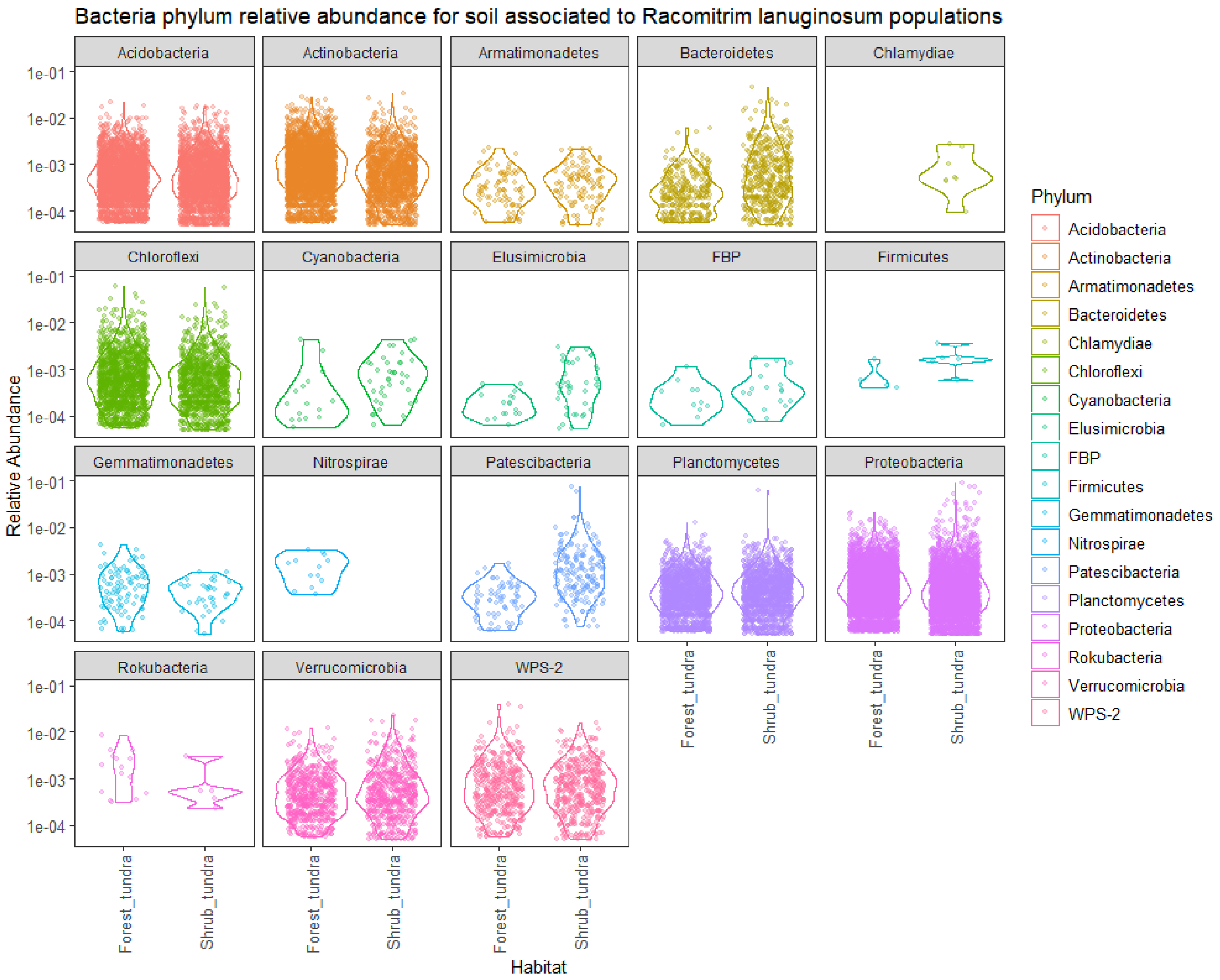


**Fig. S10** Relative abundance per phylum of bacterial ASVs of soil samples associated with two *Racomitrium lanuginosum* populations based on 16S rRNA data. The most abundant phyla are Acidobacteria, Actinobacteria, Chloroflexi and Proteobacteria. The ASV relative abundance of Nistrospirae and Chlamydiae differed between the forest and shrub tundra

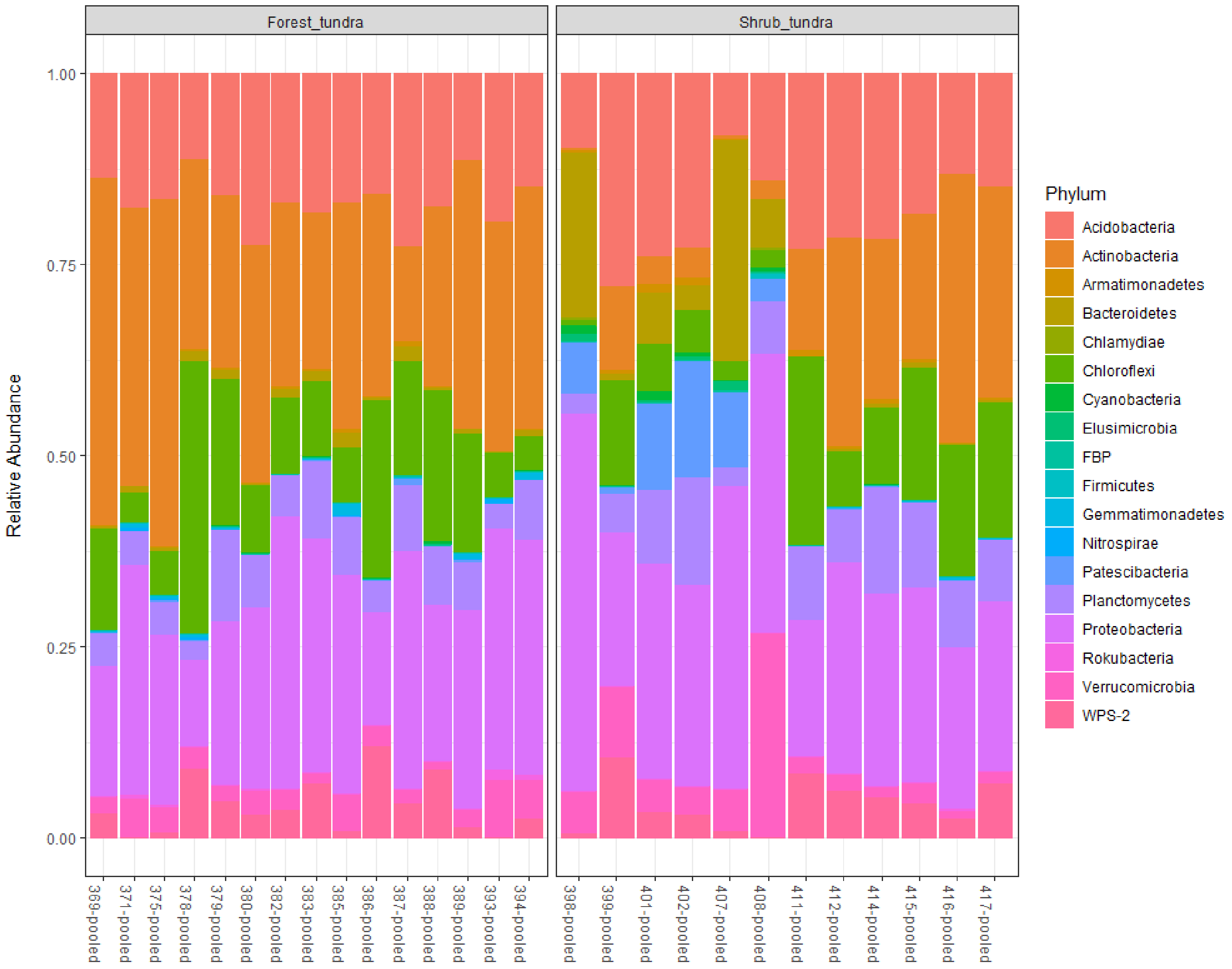


**Fig. S11** Relative abundance per phylum of bacterial ASVs of soil samples associated with two *Racomitrium lanuginosum* populations based on 16S rRNA data. The most abundant phyla are Proteobacteria, Actinobacteria and Acidobacteria. ASV abundances seem similar in forest and shrub tundra

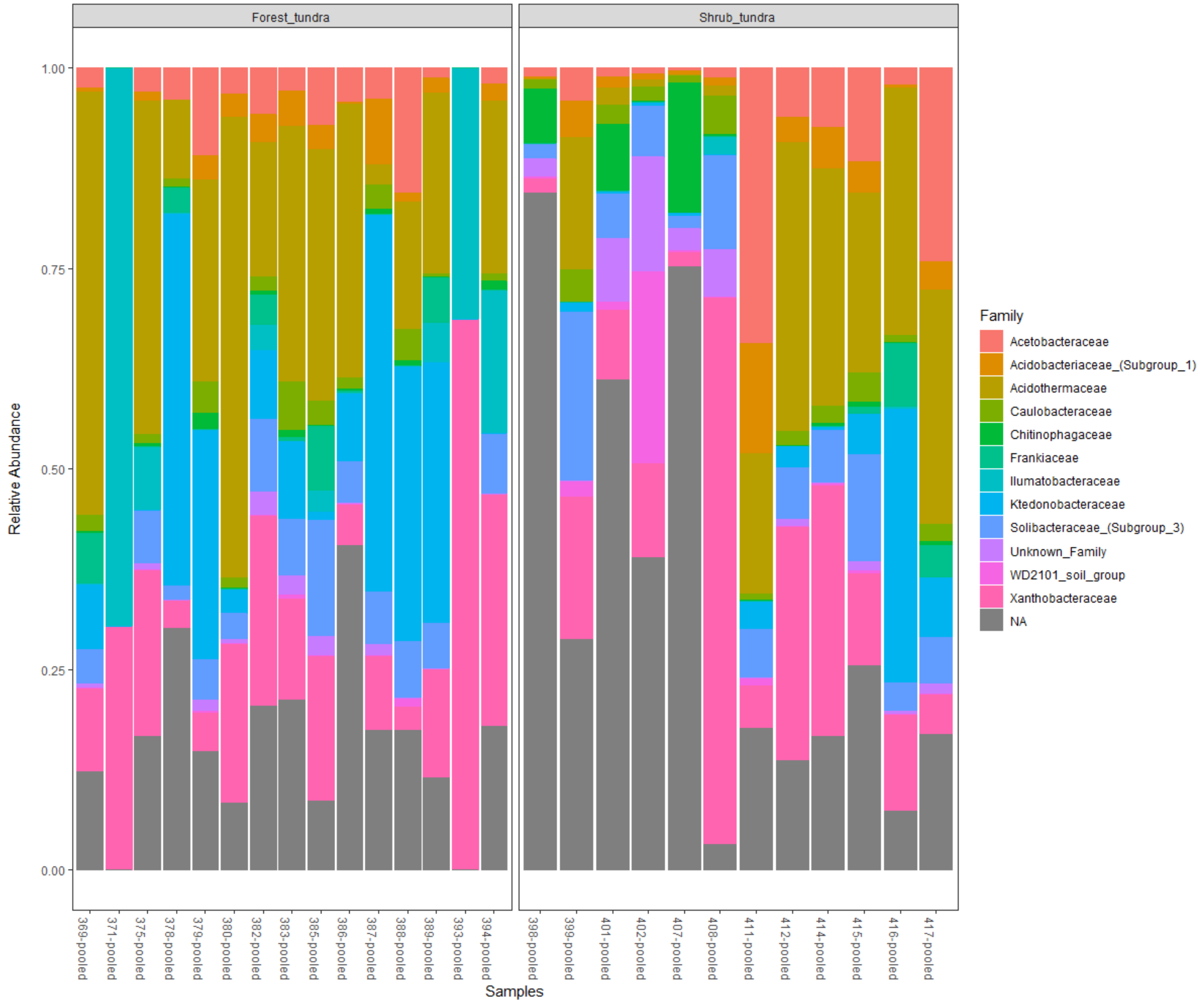


**Fig. S12** Relative abundance per family of bacterial ASVs of soil samples associated with two *Racomitrium lanuginosum* populations based on 16S rRNA data. The most abundant families are Acidothermaceae, Ktedonobacteraceae and the WD2101 soil group. Ktedonobacteracea seems to be more abundant in the shrub tundra

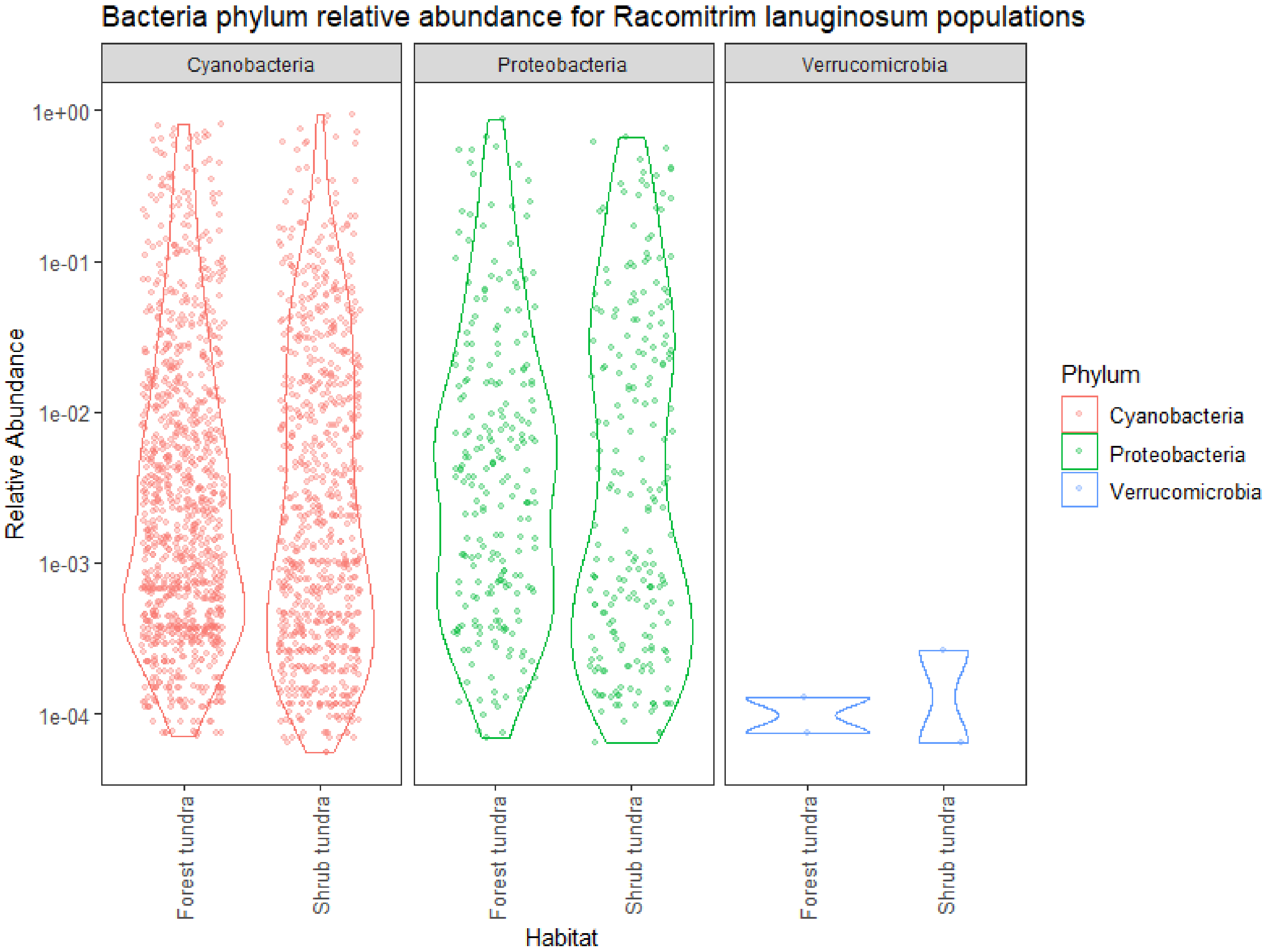


**Fig. S13** Relative abundance per phylum of diazotrophic OTUs of two *Racomitrium lanuginosum* populations based on *nif*H data. Putative diazotrophic OTU abundances seem similar between the forest tundra and shrub tundra

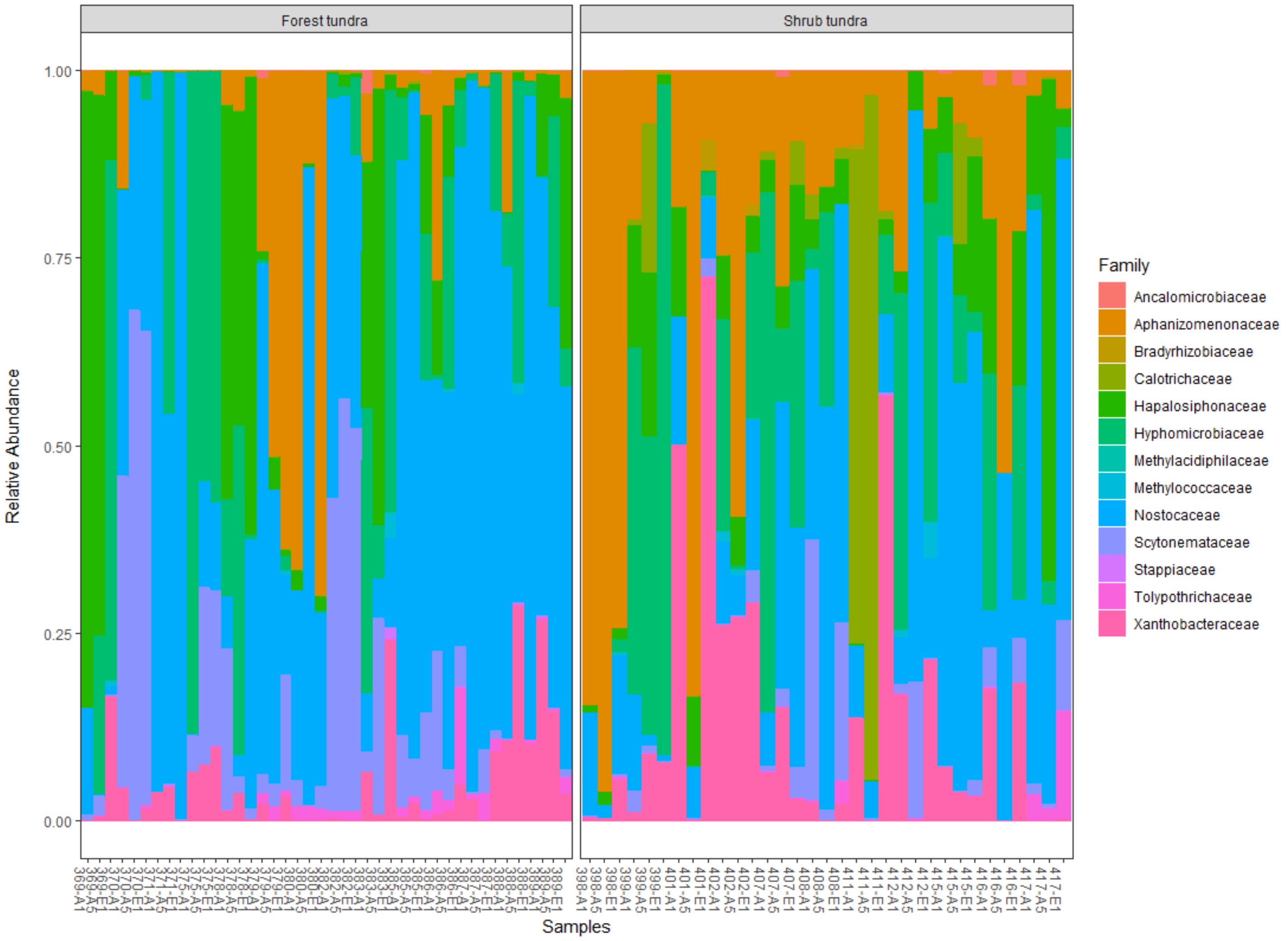


**Fig. S14** Relative abundance per family of the 20 most abundant diazotrophic OTUs of two *Racomitrium lanuginosum* populations based on *nif*H data. The Nostocaceae, Hyphomicrobiaceae and Aphanizomenonaceae families characterized dominant OTUs. Nostocaceae seems better represented in the forest-tundra

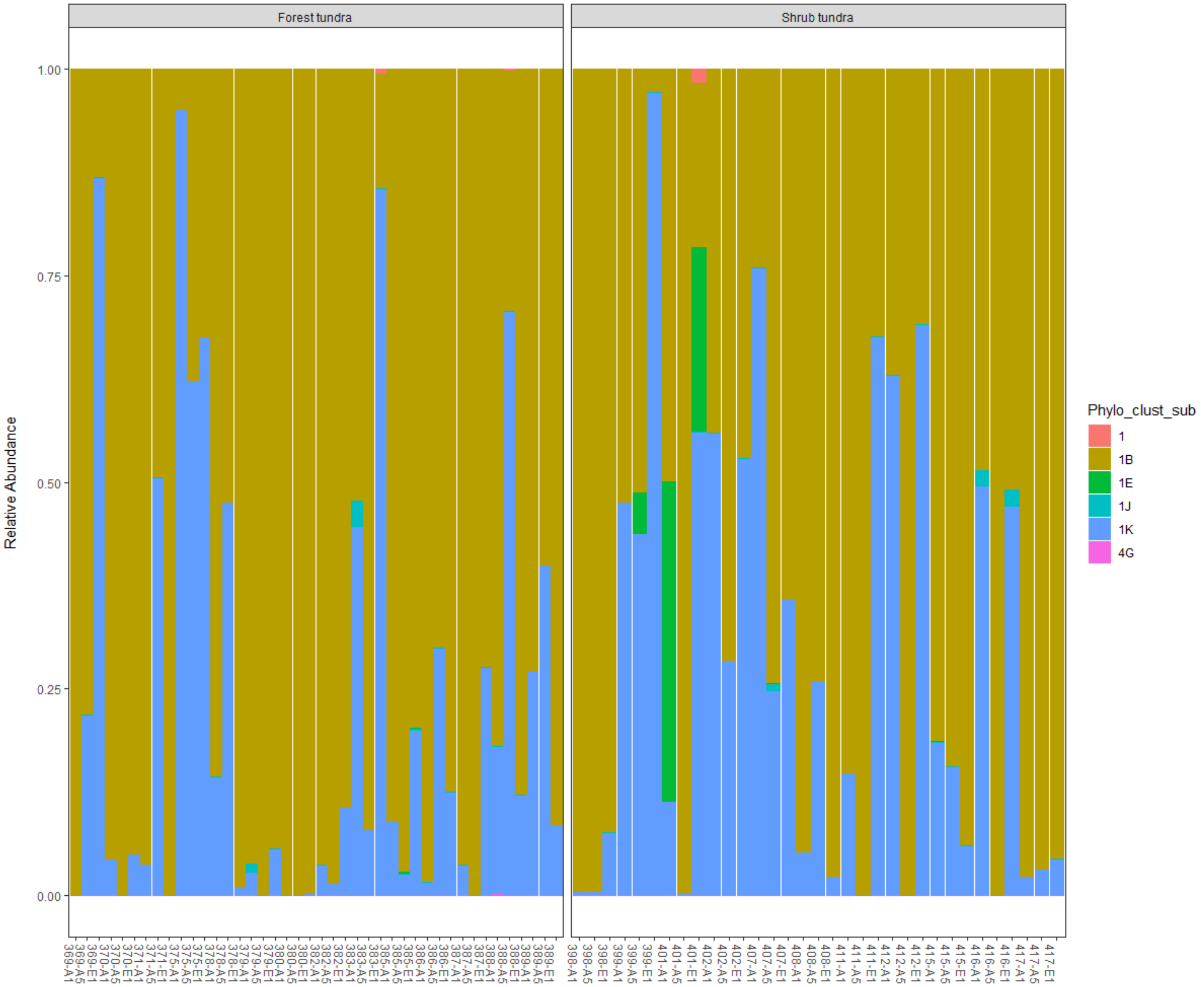


**Fig. S15** Relative abundance by phylogenetic cluster of the 20 most abundant diazotrophic OTUs of two *Racomitrium lanuginosum* populations based on *nif*H data. Most of the diazotrophic community correspond to the *nif*H phylogenetic cluster 1B, consisting of Cyanobacteria OTUs, and 1K comprising members of Alphaproteobacteria such as *Azorhizobium* and *Rhodomicrobium*

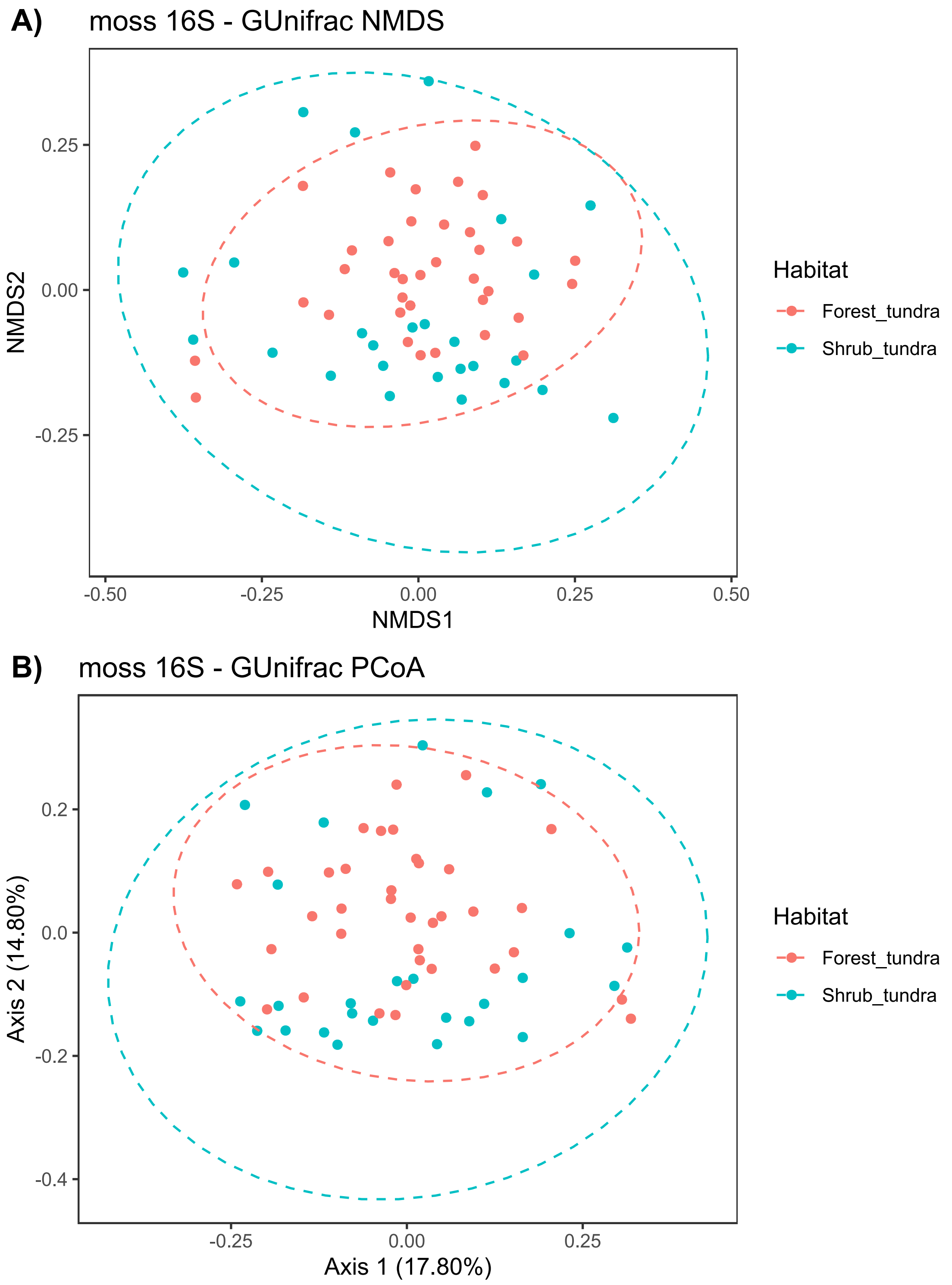


**Fig. S16** Ordination methods of *Racomitrium lanuginosum* 16S rRNA data based on GUnifrac dissimilarity matrices in two populations. A) Non-metric multidimensional scaling (NMDS) of 16S microbial data with samples color-coded according to tundra type; B) Principal coordinates analysis (PCoA) based on 16S microbial data with samples color-coded according to tundra type. The dotted circles indicate the 95% confidence intervals. Samples do not seem to cluster according to habitat

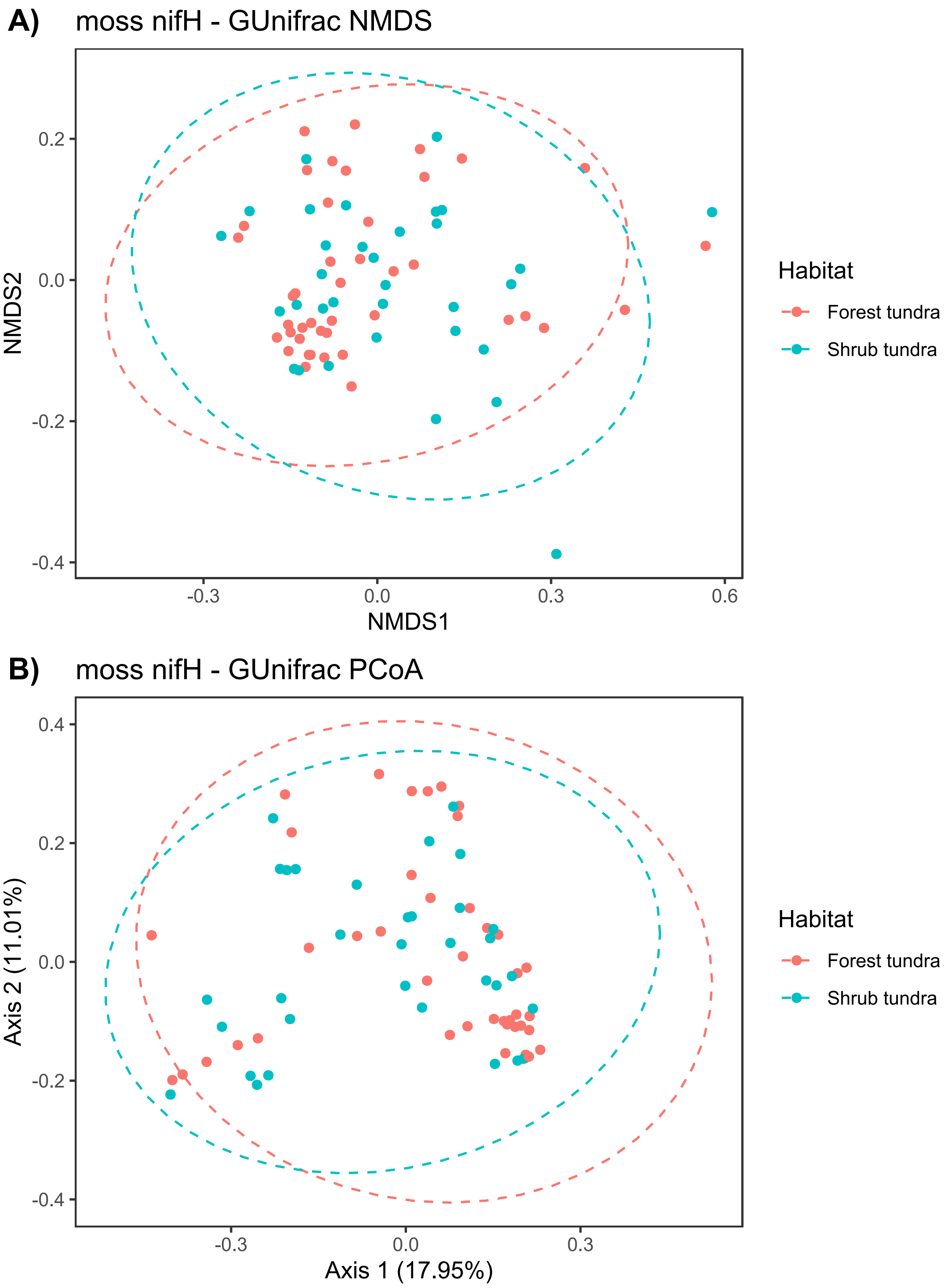


**Fig. S17** Ordination methods of *Racomitrium lanuginosum nif*H data based on GUnifrac dissimilarity matrices in two populations. A) Non-metric multidimensional scaling (NMDS) of *nif*H microbial data with samples color-coded according to tundra type; B) Principal coordinates analysis (PCoA) based on *nif*H microbial data with samples color-coded according to tundra type. The dotted circles indicate the 95% confidence intervals. Samples do not seem to cluster according to habitat

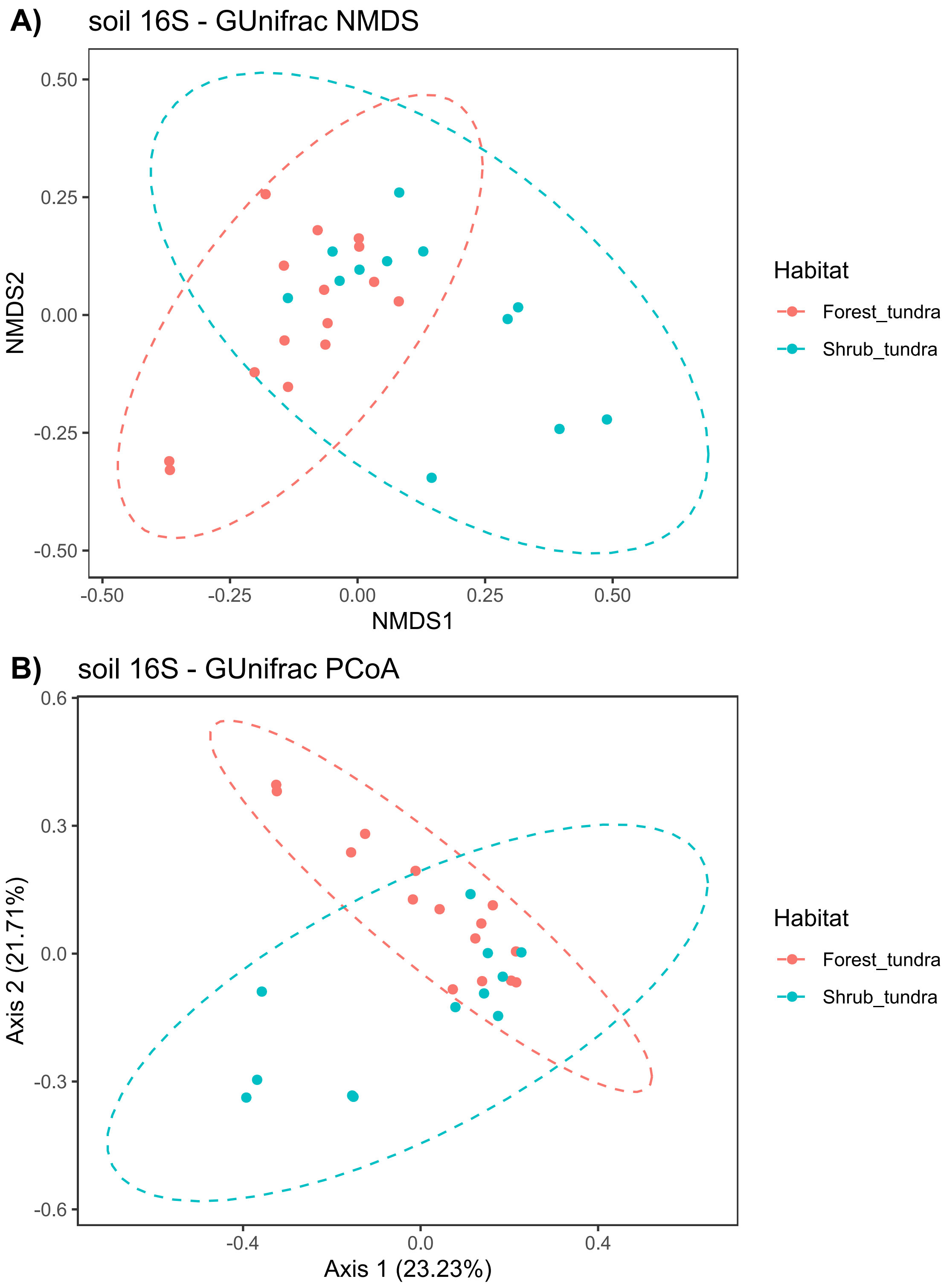


**Fig. S18** Ordination methods of soil 16S rRNA data associated with *Racomitrium lanuginosum* based on GUnifrac dissimilarity matrices in two populations. A) Non-metric multidimensional scaling (NMDS) of soil 16S rRNA microbial data with samples color-coded according to tundra type; B) Principal coordinates analysis (PCoA) based on soil 16S microbial data with samples color-coded according to tundra type. The dotted circles indicate the 95% confidence intervals. Samples seem to cluster according to habitat type

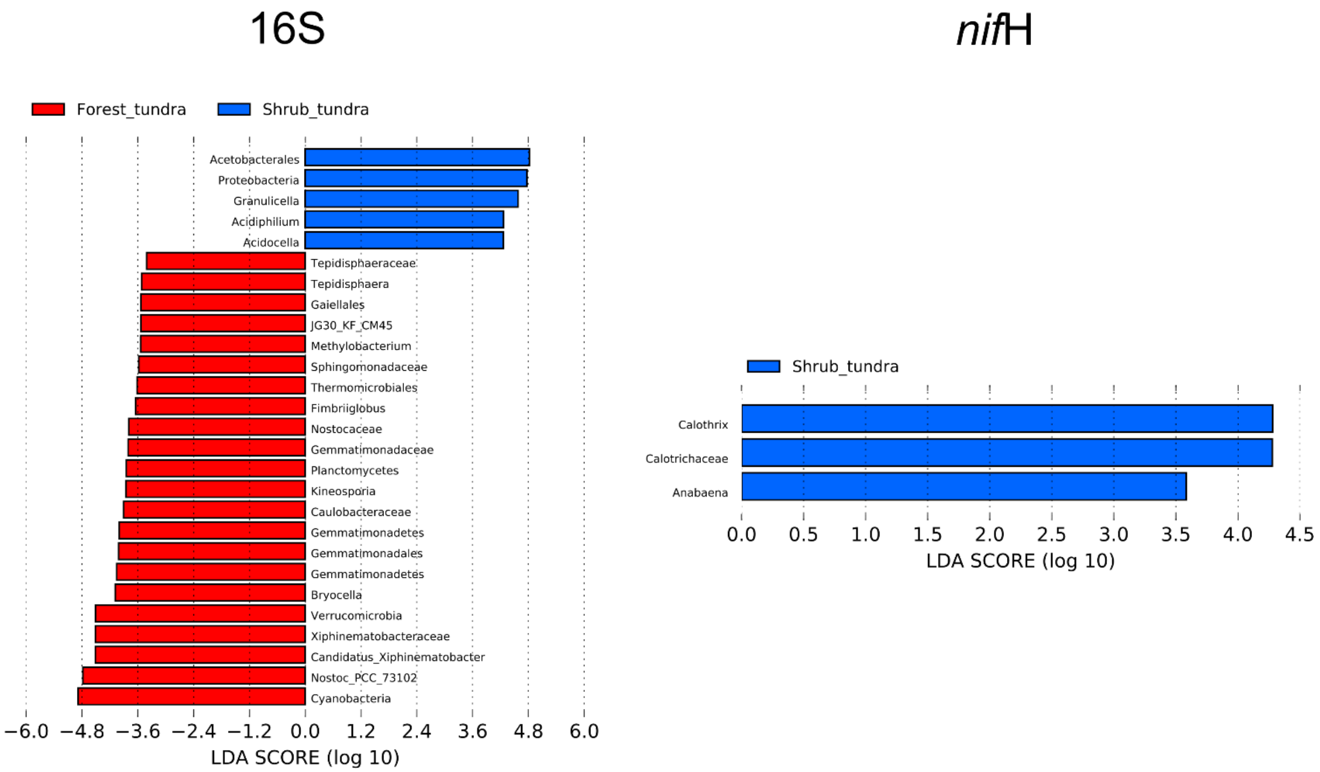


**Fig. S19** LDA scores of bacterial and diazotrophic groups based on LEfSe analyses at genus level in two tundra types. Results of the 16S rRNA LEfSe analysis are shown on the left panel. Each differentially abundant group is presented with their LDA scores, and color coded according to tundra type. The LDA scores for the *nif*H LEfSe analysis are shown on the right. Results indicate that only the shrub tundra exhibited differentially abundant diazotrophic groups at the genus level

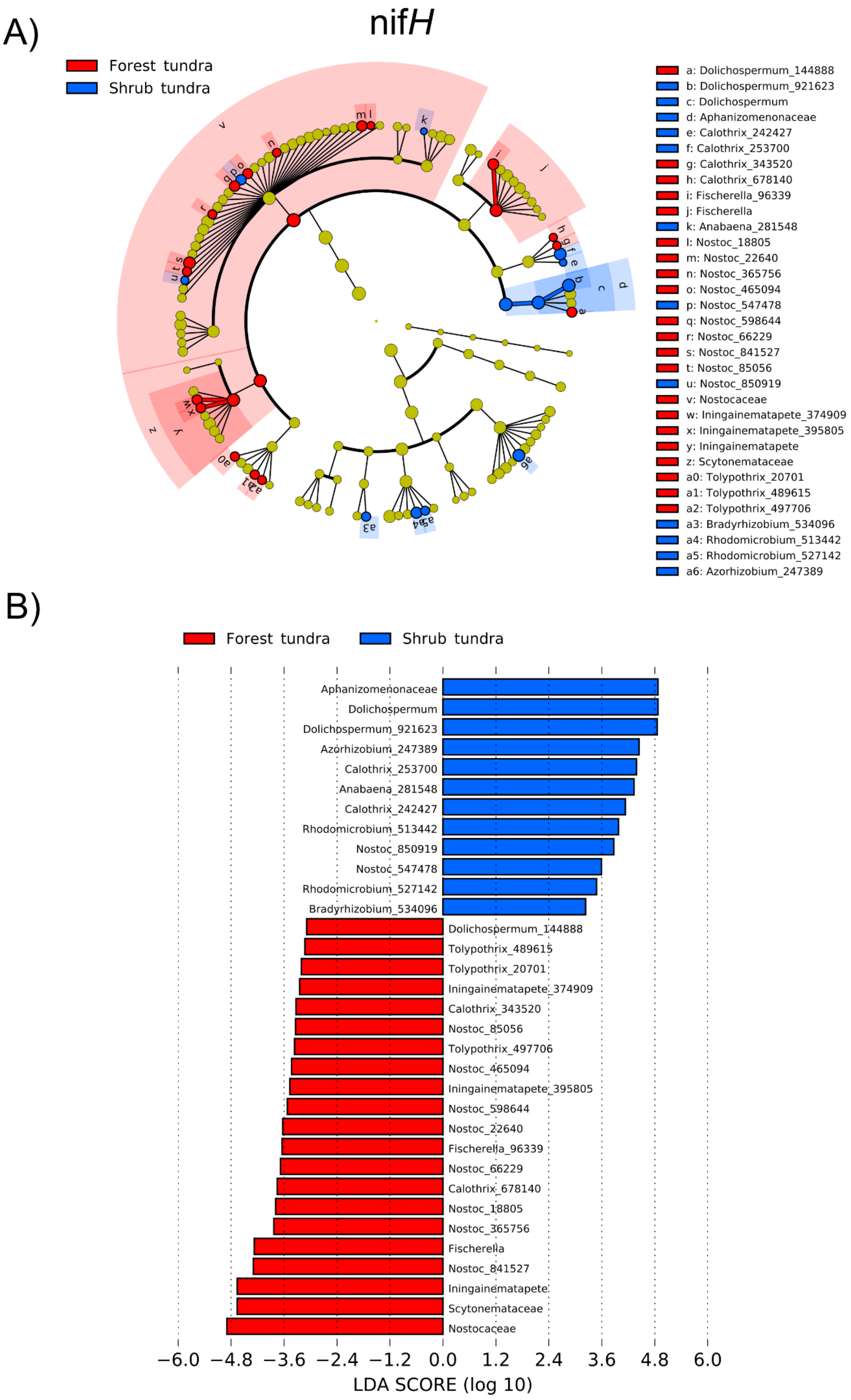
A)
nifH
Forest tundra
Shrub tundra
a: Dolichospermum_144888
b: Dolichospermum_921623
c: Dolichospermum
d: Aphanizomenonaceae
e: Calothrix_242427
f: Calothrix_253700
g: Calothrix_343520
h: Calothrix_678140
i: Fischerella_96339
j: Fischerella
k: Anabaena_281548
l: Nostoc_18805
m: Nostoc_22640
n: Nostoc_365756
o: Nostoc_465094
p: Nostoc_547478
q: Nostoc_598644
r: Nostoc_66229
s: Nostoc_841527
t: Nostoc_85056
u: Nostoc_850919
v: Nostocaceae
w: Iningainematapete_374909
x: Iningainematapete_395805
y: Iningainematapete
z: Scytonemataceae
a0: Tolypothrix_20701
a1: Tolypothrix_489615
a2: Tolypothrix_497706
a3: Bradyrhizobium_534096
a4: Rhodomicrobium_513442
a5: Rhodomicrobium_527142
a6: Azorhizobium_247389
B)
Forest tundra
Shrub tundra
Aphanizomenonaceae
Dolichospermum
Dolichospermum_921623
Azorhizobium_247389
Calothrix_253700
Anabaena_281548
Calothrix_242427
Rhodomicrobium_513442
Nostoc_850919
Nostoc_547478
Rhodomicrobium_527142
Bradyrhizobium_534096
Dolichospermum_144888
Tolypothrix_489615
Tolypothrix_20701
Iningainematapete_374909
Calothrix_343520
Nostoc_85056
Tolypothrix_497706
Nostoc_465094
Iningainematapete_395805
Nostoc_598644
Nostoc_22640
Fischerella_96339
Nostoc_66229
Calothrix_678140
Nostoc_18805
Nostoc_365756
Fischerella
Nostoc_841527
Iningainematapete
Scytonemataceae
Nostocaceae
−6.0 −4.8 −3.6 −2.4 −1.2 0.0 1.2 2.4 3.6 4.8 6.0
LDA SCORE (log 10)

**Fig. S20** LEfSe results of differentially abundant OTUs based on the *nif*H gene in two tundra types. A) Cladogram of the diazotrophic community based on *nif*H showing the differentially abundant groups for the forest tundra in red and the shrub tundra in blue. The legends indicate taxonomic groups with letters within the cladograms. B) Each differentially abundant group is presented with their LDA scores and colour coded according to tundra type. Proteobacteria diazotrophs are more abundant in the shrub tundra than in the forest tundra

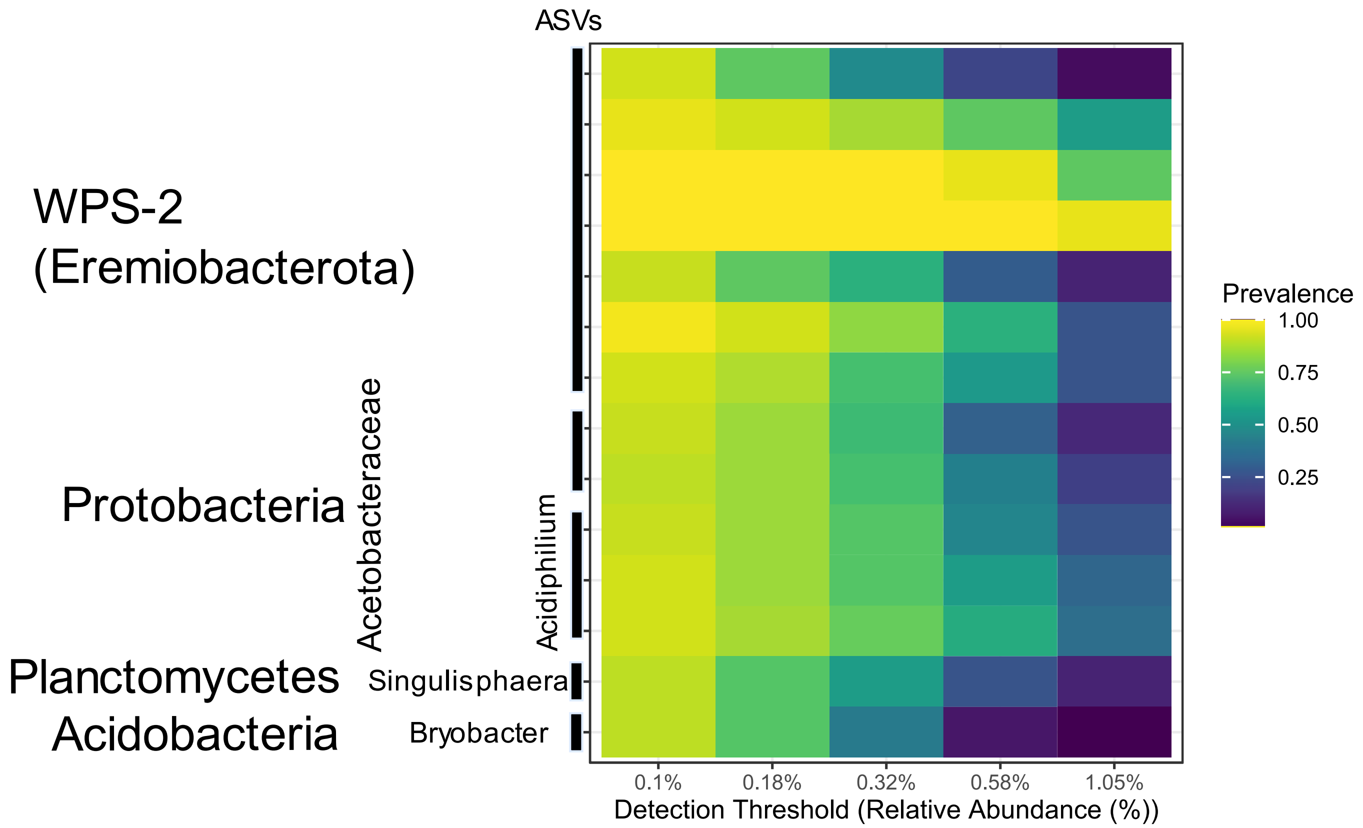


**Fig. S21** Core microbiome of *Racomitrium lanuginosum* based on ASV prevalence of the 16S rRNA gene as a response to the relative abundance detection threshold. Core ASVs are presented on the y-axis with their available taxonomic information and a prevalence higher than 0.90 for the minimum detection threshold. Detection thresholds are presented on the x-axis as the ASV relative abundance. The color scale on the left indicates the ASV prevalence according to the detection threshold ranging from minimum values on a blue scale to higher on a yellow. Increasing the detection threshold will reduce the prevalence and number of core ASVs
